# Group size influences behavioral plasticity in responses to thermoregulation-foraging trade-offs by a socially cohesive bird

**DOI:** 10.1101/2025.04.24.650232

**Authors:** David Rozen-Rechels, Danai Papageorgiou, Mina Ogino, Brendah Nyaguthii, Nina Buschhausen, Christina Hansen Wheat, James A. Klarevas-Irby, Peter Njoroge, Damien R. Farine

**Author notes:** **Corresponding author:** David Rozen-Rechels. Current address: Sorbonne Université, Université Paris Cité, Univ Paris Est Créteil, CNRS, IRD, INRAE, Institut d’écologie et des sciences de l’environnement de Paris, IEES Paris, F- 75005 Paris, France.

## Abstract

Behavioral plasticity, such as changes in habitat use and activity, can be a critical modulator of thermal pressures on endotherms. However, shifts in behaviors can meet diversified costs such as missed feeding opportunities. Individual decision-making should therefore capture the trade-off between the costs and the benefits of thermoregulation. In the case of social species, the decision process could also be facilitated or slowed down by the modulation of costs arising through the social environment. In this study, we tested how vulturine guineafowl (*Acryllium vulturinum*) change their use of open areas (where they predominately forage) according to heat using GPS data from 105 birds collected every 5 minutes for 6 months. Because animals vary in their sensitivity to risk according to group size—e.g. due to the dilution effect—we compared the responses of individuals depending on the size of the group they belong to. We also analyzed if behavioral responses translate into less precise thermoregulation by recording the body temperature of a subset of birds in two groups. We foundd that birds avoid heat by selectively using open areas and moving to cover, as well as reducing activity when temperature increases, birds use open areas less and move less. Individuals from intermediate-sized groups seemed to be able to use open areas during warmer conditions compared to individuals from small and large groups (10% higher probability of use). However, active birds in the open did not present hyperthermia, suggesting that behavioral changes are efficient or that individuals have other efficient strategies such as physiological cooling. Together, these results indicated that responses to high temperatures are complex, as they included not only a range of environmental constraints but that responses can also vary according to social context.

## Introduction

Climate change is predicted to not only drive an increase in average temperature but also to increase the frequency and the intensity of the occurrence of extremely high temperatures (IPCC 2021). High temperatures can strongly constrain the physiology of organisms, ultimately leading to lethal hyperthermia and dehydration if temperatures exceed the organism’s performance range (J. B. Williams and Tieleman 2005; Angilletta 2009; McKechnie and Wolf 2010). Animals can, however, use short-term behavioral adjustments which buffer the effect of high temperatures on physiology (Marais and Chown 2008; Huey et al. 2012; Long et al. 2014; Sunday et al. 2014).

This behavioral plasticity, often called “behavioral buffering”, can range from postural changes limiting the exposure of the body to hot surfaces or solar radiation, to changes in habitat use and activity patterns (Cunningham, Martin, and Hockey 2015). Typically, the heterogeneity of the landscape—featuring a mosaic of sunny hot micro-habitats and shaded shelter—can allow animals to avoid high temperatures and facilitate thermoregulation by selecting microhabitats close to their physiological thermal preferences (Long et al. 2014; Sears et al. 2016). This flexibility is likely to make behavior particularly critical for ectotherm thermoregulation (Bauwens, Hertz, and Castilla 1996; Angilletta 2009). It is also a modulator of body temperature in endotherms by reducing water losses through evaporative cooling (Wolf and Walsberg 1996; Cain III et al. 2006; Hetem et al. 2012; Smit et al. 2016; Oswald et al. 2019). Thus, in principle animals can maintain their body temperature constant in an optimal range through behavior when the environment allows them to.

Behavioral thermoregulation is, however, energetically costly (Angilletta 2009). In addition to energetic expenditure in movement, shifts in activity patterns and micro-habitat use in response to heat can lead to “missed-opportunity” costs, such as a suboptimal use of food or water resources, an increased exposure to predators, or a reduction in the expression of reproductive or social behaviors—especially when behavioral decisions are mutually exclusive (du Plessis et al. 2012; Cunningham, Gardner, and Martin 2021; Stofberg et al. 2022). When faced with such thermoregulation trades-offs, we expect behavioral decisions to maximize the overall benefits and minimize overall costs of the constraints of the environment. As a consequence, a behavior might appear to be suboptimal when studied under the frame of thermoregulation alone.

Examples of this are large savanna grazers that avoid foraging in cooler periods of the day when faced with higher predation pressure (Veldhuis et al. 2020) and lizards that shift their micro- habitat preferences to cooler and moister places under water restriction at the cost of taking advantage of better thermal conditions for thermoregulation (Rozen-Rechels et al. 2019; 2020). But, in other cases, heat avoidance must be prioritized. For example, southern yellow-billed hornbills (*Tockus leucomelas*) and Alpine ibex (*Capra ibex)* engage in thermorgulation at the expense of foraging and body condition (Mason et al. 2017; van de Ven, McKechnie, and Cunningham 2019). Therefore, behavioral adjustments for thermoregulatory needs under heat stress are not trivial to predict, especially when those implies a trade-off with meeting foraging needs.

If heat avoidance is prioritized in the animal’s decision-making, we expect body temperature to remain constant and close to the physiological optimum. However, the costs associated with water losses under high temperatures might enhance different physiological responses. Water conservation behaviors for example would reduce dehydration at the expense of suboptimal body temperatures (Anderson and Andrade 2017; Rozen-Rechels et al. 2019; 2020). This is not only true in ectotherms but also in endotherms, if the latter can relax the need to maintain a precise body temperature under thermal conditions that challenge their water and energetic balances (Wooden and Walsberg 2002; Hetem et al. 2016; McKechnie and Wolf 2019). Some arid habitat birds could even tolerate a facultative hyperthermia, thus limiting dehydration (Gerson et al. 2019). The same way as ectotherms’ behavioral strategies could be studied indirectly through the analyses of individuals body temperatures (Blouin-Demers and Nadeau 2005; Rozen-Rechels et al. 2021), the efficiency of endotherms’ behavioral decision facing heat could also be understood by studying the variations of their body temperatures (Thompson, Cunningham, and McKechnie 2018).

Until recently, behavioral responses to heat were predominately studied in solitary individuals. While this is relevant in many species, it neglects that social interactions are critical in movement and habitat use decision-making processes (Couzin et al. 2005; Strandburg-Peshkin et al. 2015; Hansen et al. 2024). Conspecifics can have a substantial influence on foraging behaviors.

Individuals in groups benefit from information sharing, which can not only improve access to resources such as food but also thermal refuges in heterogeneous environments (Dall et al. 2005; Cantor, Aplin, and Farine 2020). Group living also dilutes predation pressures and allows the emergence of collective antipredator strategies such as sentinel behaviors and mobbing (Kenward 1978; Graw and Manser 2007; Lehtonen and Jaatinen 2016). We might then expect food, temperature and predation constraints to synergistically favor larger groups if these can better offset trade-offs with predation. However, larger groups also tend to slow down the decision-making process and reduce the group movement speed due to coordination challenges (Herbert-Read et al. 2013; Strandburg-Peshkin et al. 2015; 2017; Papageorgiou and Farine 2020b; Klarevas-Irby, Nyaguthii, and Farine 2025), potentially limiting their ability to take advantage of heterogeneous landscape features. The trade-off between group advantages in information sharing and protection, and the constraints in movement, might then favor an intermediate—or ‘optimal’—group size. However, group size effects on micro-habitat selection and behavioral buffering of heat stress have not been explored.

In this study, we investigated the interplay between foraging and heat constraints on the habitat use of a large social bird, the vulturine guineafowl *Acryllium vulturinum* (Hardwicke 1834).

Given their relatively large body size (1.5–1.8 kg), we expected heat dissipation to be challenging for vulturine guineafowl and, thus, that they should rely on behavior for thermoregulation (Smit et al. 2016; Pattinson et al. 2020) as has been shown in the closely related helmeted guineafowl *Numida meleagris* (Rakowski et al. 2019). However, thermoregulation is likely to be traded off against foraging, the latter being predominately done in open grassy areas exposed to the sun and rich nutrients from being used overnight by large herbivores (Young, Patridge, and Macrae 1995). We used large-scale and long-term data from GPS trackers fitted to approximately 10% of a population of wild vulturine guineafowl to test how birds prioritize between different measures of heat constraints when deciding to use these foraging areas. We also tested whether individuals shift their habitat use differently according to the size of their group, balancing the complexity of decision-making with the anti-predation and collective intelligence benefits that larger groups experience.

As a result of the trade-off with the ability to effect rapid decision-making, we expected that individuals living in large groups may be more risk-averse in terms of temperature. We also expected that groups of intermediate size will spend more time in the open under heat relative to individuals in small or large groups. Finally, given that vulturine guineafowl are endemic to the arid savannahs of east Africa, we expected to observe birds managing trade-offs under higher heat conditions by exhibiting hyperthermia, which we measure using internal body temperature loggers on a subset of GPS-tracked birds. We lay out all of our predictions in Table 1.

**Table 1.** Hypotheses and predictions on the priorities of the vulturine guineafowl between heat and food avoidance and foraging which are compared in the study. VGFs: vulturine guineafowl. T_sun_: operative temperature in the sun. T_diff_: difference of operative temperature the sun and the shade.

| # | Hypothesis | Predictions | Model formula |
| --- | --- | --- | --- |
| 1 | Random foraging behavior | VGFs select randomly the habitat to use | Habitat use index $\sim 1$ |
| 2 | Temperature-driven foraging behavior | VGFs use open habitats less with increasing sun temperature in order to avoid hyperthermia | Habitat use index $\sim T_{sun}$ |
| 3 | | VGFs use open habitats the most at intermediate sun temperature in order to avoid hyperthermia but also to take advantage of warmer temperatures at the start of the day | Habitat use index $\sim T_{sun} + T_{sun}^2$ |
| 4 | | VGFs use open habitats the most when the temperatures in open and cover habitats are similar, i.e. when heat constraints are the lowest or when there is no increased thermal cost to go in the sun compared to staying in the shade | Habitat use index $\sim T_{diff}$ |
| 5 | | VGFs use open habitats the most when the temperature in the open is higher than under cover when heat constraints are low (early or late in the day, cloudy days), but avoid open habitats when the temperature gets too high while staying in the shade allow cooling at reduced costs | Habitat use index $\sim T_{diff} + T_{diff}^2$ |
| 6 | Group foraging behavior | The use of open habitats is maximal at medium group size as the benefits of predation risk dilution in large groups and low competition in small groups trade-off | Habitat use index $\sim Group\ size$ |
| 7 | Temperature-driven group foraging behavior | Prediction #2 + Medium-sized groups use open habitats at higher sun temperatures than other groups | Habitat use index $\sim T_{sun} \times Group\ size$ |
| 8 | | Prediction #3 + Medium-sized groups use open habitats at higher sun temperatures than other groups | Habitat use index $\sim [T_{sun} + T_{sun}^2] \times Group\ size$ |
| 9 | | Prediction #4 + Medium-sized groups use open habitats at higher sun temperatures than other groups | Habitat use index $\sim T_{diff} \times Group\ size$ |
| 10 | | Prediction #5 + Medium-sized groups use open habitats at higher sun temperatures than other groups | Habitat use index $\sim [T_{diff} + T_{diff}^2] \times Group\ size$ |

## Material and Methods

### Study system

Vulturine guineafowl (hereafter called VGFs) are predominately terrestrial birds that live in highly cohesive social groups ranging from a dozen to over 60 individuals (Papageorgiou et al. 2019). Group membership is stable, but groups can respond to ecological conditions by temporarily splitting up (to breed; Nyaguthii et al. 2025) or by temporarily associating with other groups (during dry periods, Ogino, Strauss, and Farine 2023). Herein our references to ‘group’ correspond to the stable social groups that individuals spend most of their lives in (Farine *in review*), which reflects a stable social unit at the intermediate level of their multilevel society. In this paper, group size refers to the number of conspecifics that an individual is currently with (which can include individuals from other stable groups). Groups follow shared decision-making processes to decide where to move, meaning that rather than following a dominant leader, all individuals can contribute to the group’s next actions (Papageorgiou and Farine 2020a; Papageorgiou, Nyaguthii, and Farine 2024). Their diet is composed of grass root buds, seeds and small arthropods that the birds mainly find on glades, i.e. open areas rich in nutrients (Young, Patridge, and Macrae 1995), showing a higher propensity to forage and greater foraging effort on these open areas (Appendix 1).

Our study site is a c. 12km^2^ dry woody-savanna in the south of Mpala Research Center (MRC), in Laikipia, Kenya. This tropical ecosystem is typically described by two main wet seasons (October to November and from April to June) separated by dry seasons characterized by low rainfall and little to no green vegetation. VGFs home range size as well as the distance moved every day drastically increase during the dry seasons when resources are more clumped into open areas (Papageorgiou et al. 2021). The current study focuses on an extraordinarily long dry period that lasted from December 2020 to July 2021 (Appendix 2: Figure S2).

VGFs are a prey species that faces pressure from both aerial (raptors such as martial eagles *Polemaetus bellicosus*, Heine 1890; tawny eagles *Aquila rapax*, Temminck 1828; and African hawk-eagles *Aquila spilogaster*, Bonaparte 1850) and terrestrial (e.g. jackals, hyenas) predators. While encounters with opportunistic terrestrial predators occur supposedly at random (Cooper, Holekamp, and Smale 1999; Moehlman 2014), i.e. indiscriminately in open and covered patches, VGFs are particularly visible (and sensitive) to soaring predators (raptors) in open areas.

### GPS tracking data

From early February 2021 to the end of July 2021, high-resolution solar-powered GPS tags (15g Bird Solar, e-obs Digital Telemetry, Grünwald, Germany) were fitted to 118 different individuals (from both sexes and all adults) across 20 identified social groups. In order to prevent feathers from covering the tags’ solar panel, loggers were elevated using neoprene pads. The tag, Teflon harness, and the platform used to raise the tag above the feathers together weighted c. 20.5g, less than 2% of the weight of the bird (below the 3% recommendation for animal welfare, see Bodey et al. 2018; Portugal and White 2018 for further discussion). The birds tracked in this study were tagged between October 2016 and June 2021. Data were downloaded remotely every 2 days using a BaseStation II (e-obs Digital Telemetry, Grünwald, Germany) and uploaded on the Movebank repository (Kays et al. 2022). The majority of the birds in the study population (including all GPS-tagged birds) were fitted with a unique combination of colour leg bands, allowing identification in the field.

We excluded data collected on the day of deployment for birds that joined the study along the way, as well as data collected during periods when a bird’s group was being trapped. We also excluded birds living in the group habituated to human living in the research center as they use a fenced, highly vegetated area with different predation pressure and resource availability, as well as subadult birds which were tagged as part of a study on dispersal (Klarevas-Irby, Wikelski, and Farine 2021). We collected individual locations from 07:00 to 18:00 (EAT; when the birds are out of the roosts), with a burst of 10 locations (1Hz) every 5 minutes. We identified and excluded outliers’ location when two consecutive locations were farther than 600m (this threshold have been defined by visualizing the tracks and observing the distribution of consecutive locations distance). In this project, we only kept the 10^th^ location of each burst which is supposed to be the most accurate one (He et al. 2023), thus down-sampling our tracking data to one measure every 5 minutes. Finally, we removed birds with fewer than 500 total locations (3 birds). Our final dataset consisted of movement trajectories from 105 different birds.

### Community composition and group size

Birds were censused daily by driving along a network of vehicle tracks crossing the study area. When encountering a cohesive set of individuals, we recorded the number of individuals and identified all the marked birds present (or until birds moved into vegetation). Membership can be dynamic during dry periods—such as in our study period—as the long-term stable groups can temporarily merge as part of the multilevel society. Thus, we estimated the group size experienced by each tagged individual each month using a network community detection approach as described in Ogino, Strauss, and Farine (2023). Specifically, we extracted in census data all observations in which tagged birds were identified each month. For each observation, we compared the birds identified to the network community membership (based on all the data, per Ogino et al. 2023) and counted (1) the number of observed birds that belong to the community that month, (2) the number of observed birds that do not belong to the community that month and (3) the number of known birds in the community based on community detection. We calculated the ratio (1) / [(2) + (3)] and defined each individual’s group size based on the number of individuals identified in their observation with the highest ratio. If two group observations showed the best ratio, we used the observation with the maximum number of birds. If an individual had not been censused during a given month, we predicted its group size using linear interpolation from the month before to the month after. If it had not been observed the month before or after, we considered group size to be the same as the one counted when the data was available. We then defined “small groups” for a given month to be those with group sizes in the lowest tercile (average: 34 ± 9 SD individuals, from 16 to 47 individuals), “big groups” to be those with group sizes in the upper tercile (average: 74 ± 13 SD individuals, from 58 to 109 individuals), and “median groups” those with intermediate group sizes (average: 52 ± 3 SD individuals, from 48 to 57 individuals).

### Body temperature

Between February 25^th^ and March 1^st^ 2021, 23 VGFs from two groups were implanted with an ECG-logger (ECG-tag 1AA2, e-obs Digital Telemetry, Grünwald, Germany; 25g; 38 x 23 x 19mm) in addition to GPS tags (the sum of the weights of the loggers ranged from 2.6 to 3.4% of the weight of the birds). ECG-loggers record heart-rate data (not used in this study, see Brandl et al. 2025) and internal body temperature with an inaccuracy of maximum 0.5°C. The ECG logger was implanted in the thoraco-abdominal cavity of anesthetized birds. The logger was fixed to the abdominal wall with an absorbable suture (Monosyn® 4/0, B. Braun AG, Melsungen, Germany), with the longer electrode placed close to the heart. All implantation details are described in Brandl et al. (2025) Data were downloaded remotely every fourth night. We excluded 5 birds that were predated or died in the month following implantation. Five loggers that showed unexpected body temperature records below 34°C or higher than 47°C were also excluded from the analysis. Body temperature was sometimes recorded once every 5 minutes or once every 20 seconds. We homogenized the dataset by averaging body temperature every 5 minutes in the second case, hereafter called *T_body_* (103679 occurrences for 16 birds, Appendix 3: Figure S1). We then matched *T_body_* with recorded locations of 14 birds (66399 locations as some birds’ locations were only recorded once every 4 days and the locations of one bird were not recorded during the study due to a faulty GPS tag). For each individual we then calculated the resting body temperature which we defined as the average body temperature of the individual when it is resting in the shade across the whole study period. A bird was defined as inactive when its speed (here distance moved between two locations divided by the location) was below 0.01 m/s (i.e. less than 3m moved in 5 minutes, Appendix 3: Figure S2) and it was located under cover. We did not use instant speed as body temperature would not change as fast as a bird would change its behavior. This measure of activity is thus an integration of what the bird was doing in the last 5 minutes. For active birds, we then calculated the deviation between *T_body_* and the average resting body temperature per individual, hereafter called the body temperature anomaly Δ*T_body_*.

### Remote sensing and landscape description

We classified habitat within the home range of our study population as open (where VGFs can find food but where they are also subject to both aerial predation pressure and higher thermal constraints) and closed (with lower food availability but also lower predation and thermal constraints). To do so, we used the ESA Sentinel-2 multi-spectral images to calculate vegetation presence metrics such as NDVI (Papageorgiou et al. 2021). We downloaded every 10m-resolution satellite image of MRC (tile 37N BA) since they were recorded and made available (October 2017) until the end of this project (early August 2021) and cropped it to restrict the analysis to the zone used by the individuals of our study. Some additional parts of the landscape, which contain black cotton soils, were also removed because of darker soil color made habitat classification less accurate (Appendix 2: Figure S1); the parts removed contained 15100 GPS locations of 20 different individuals (i.e. 1.4% of the dataset). Clouds were cropped from the image and images with more than 20% cloud cover were removed from the time series. We calculated NDVI raster maps (Normalized Difference Vegetation Index, eqn. 1), NDWI maps (Normalized Difference Water Index, water content in vegetation, eqn. 2, Gao 1996) and brightness maps (eqn. 3, Valero et al. 2016) maps for each satellite image.

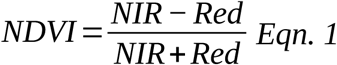

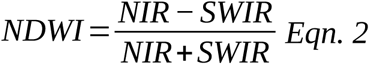

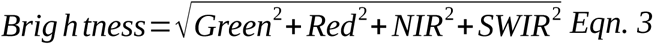

with NIR meaning Near InfraRed, SWIR meaning Short-Wave InfraRed. We then summarized these indices from October 2017 to the beginning of August 2021 into average, standard deviation, minimum, maximum and median raster maps. We then performed a simple Random Forest procedure to classify each pixel into either open or closed habitat (Breiman 2001).

We observed that the landscape of our study gives access to open and cover zones all over its area at the VGF movement scale, i.e. all groups can equally access both habitats wherever they are (Appendix 2: Figure S4 and S5). We also calculated the proportion of open pixels in a buffer of 120m-radius (99%-quantile of the distances walked by birds in 5 minutes) centered on each focal pixel, hereafter called “open patch availability”. This value allowed us to account for open vs. cover availability around each bird location in statistical models. The whole procedure is detailed in Appendix 2 and code is provided in Zenodo (the url will be provided upon publication).

### Operative temperature variations

To approach the thermal constraints experienced by VGFs in cover and in the open, we measured proxies of operative temperature (Dzialowski 2005). “Operative temperatures” are measures of the temperature of an object with the same thermal properties as the target organism, integrating heat exchanges such as conduction, convection, solar radiation, and wind cooling, thus capturing the body temperature of the organism without thermoregulation. Black copper spheres allow the measurement of relatively good proxies of operative temperatures (Walsberg and Weathers 1986) despite several issues making them unsuitable for precise thermal physiology studies (Bakken and Angilletta 2014). As our measurement errors were homogeneously distributed (i.e. because we made contrasts between microhabitats) these should not have impacted our conclusions.

Technical limitations with materials available at the field site led us to build hollow black iron cylinders which we painted with white dots to mimic VGFs (*ca* 20x20cm, hereafter called “operative models”, Figure 1A). Operative models were made of two parts: the lower one was welded to a *ca* 35cm iron bar which will be inserted 15cm deep in the soil in order to maintain the model *ca* 20cm above it such as a VGF body. We stretched and fixed a gauze across the center of each cylinder on which we mounted a DS1922L iButton (Maxim Integrated) temperature logger programmed to record temperature every 5 minutes. We deployed 20 operative models across the landscape used by the birds, during the whole duration of the study, set up in pairs space a few meters apart such that one model was in the sun and the other in the shade (Figure 1A). iButtons were replaced and downloaded every two weeks. Damaged operative models (e.g. by elephants) were repaired or replaced throughout the study and data collected by these models were excluded from the last time when it had been observed intact to the time of replacement. Data were visually monitored in order to identify operative models which could have been in the wrong micro-climatic conditions at some hours of the day (in the sun instead of shade for example). In this case, data were excluded throughout the period when the operative model was not at the right place (this could change during the course of the study due to changes in the sun course). We then calculated time series of average operative temperature in the sun (*T_sun_*, Figure 1B and Appendix 4: Figure S1) and in the shade as well as time series of the difference of average operative temperature in the sun and in the shade (*T_diff_*) every 5 minutes. Each bird location was then associated to an instant and concomitant *T_sun_* and *T_diff_*averaged for all operative temperature models. We used instant temperature conditions (instead of hourly or other type of averaged temperature conditions) to predict shifts between open and cover habitats in this study to detect possible responses to short-term drop in temperature as the consequence transient cloud cover. We also calculated the average *T_sun_* and *T_diff_* in the last 15 minutes and in the last hour before each location record to test the robustness of our results to the temporal scale of thermal conditions variables.

**Figure 1.**
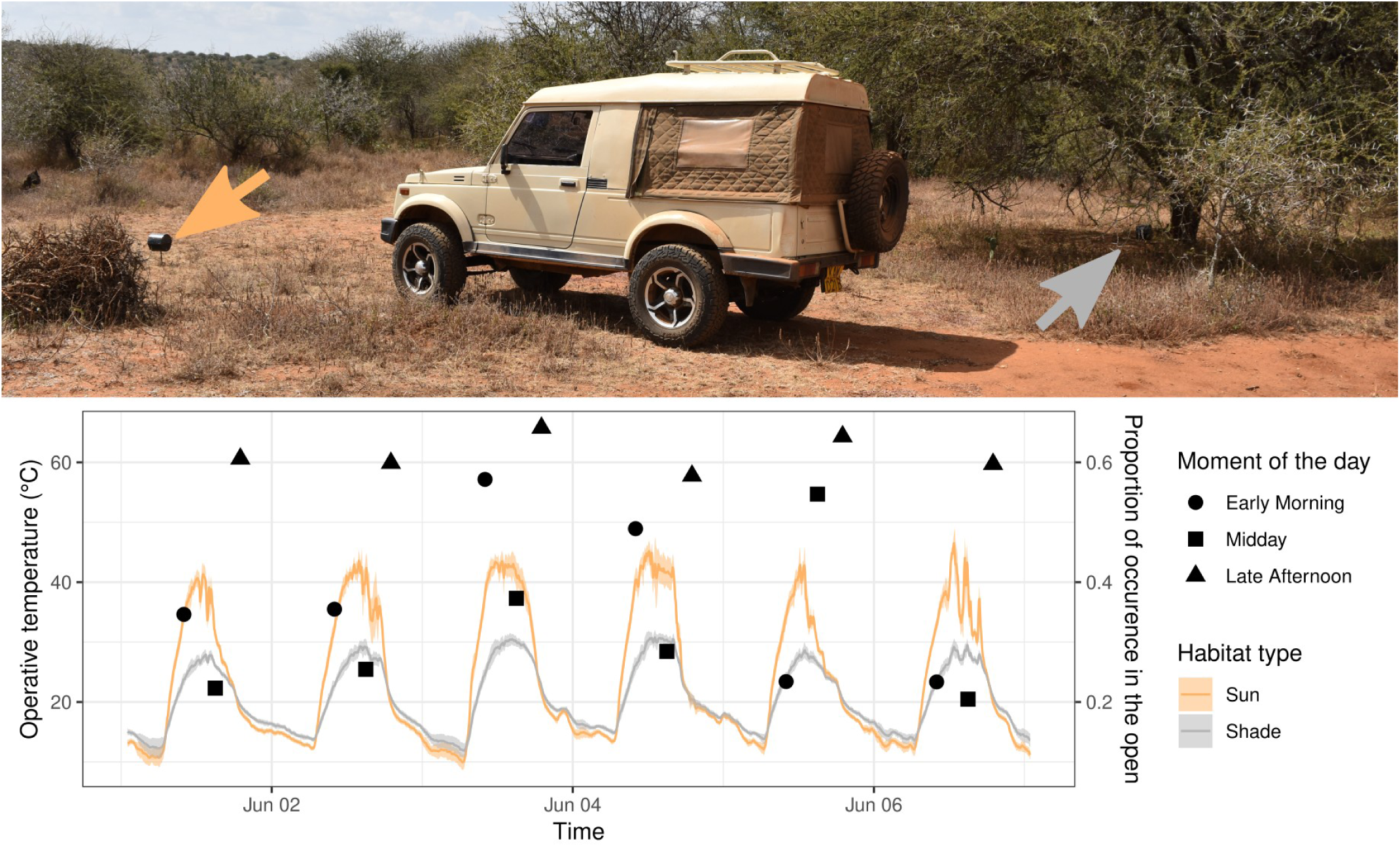
Operative temperature models. The orange arrow shows the model in open habitat and the gray arrow showing the model in cover. The car is provided for scale. The plot shows the variations of the operative temperature in the open averaged for all models in the open (i.e. *T_sun_*) and operative temperature in cover averaged for all models in cover, for a subset of dates (for clarity). Ribbons show the standard deviation. Points are showing the proportion of occurrence in the open of VGFs for 3 separated moment of the day indicated by shape: early in the morning (circles, 07:00 to 08:59 EAT), midday (squares, 12:00 to 13:59 EAT), and late afternoon (triangles, 16:00 to 17:59 EAT).

### Statistical analyses

All statistical analyses were conducted in R (4.2.1 “Funny-Looking Kid”). Generalized linear mixed models (LMM or GLMM) were fitted with the “glmmTMB” library. Models in model comparison procedures were fitted using a Maximum Likelihood (ML) estimation. Final models were analyzed after being fitted with a Restricted Maximum Likelihood (REML) estimation. Residuals distribution and homoscedasticity as well as the leverage effects of outliers or overdispersion of data were assessed in the selected (final) models with the “DHARMa” library. If we detected a significant outliers leverage effect, we tested the robustness of our results after outlier removal. If we detected a significant overdispersion of data in binomial error models, we added an observation-level random effect and tested the robustness of our results after correction. In both outliers and overdispersion case, if the problem was not solved after correction but results remained qualitatively the same, we considered our model to be robust. We visually analyzed QQ- plots considering that the very large size of our data would always lead to significant difference between the residuals distribution and a Gaussian distribution.

### Variations of habitat use

We analyzed which prediction (see Table 1) best explained the probability of being in the open (i.e. under sun; 1,018,675 locations, 9909 individual.day trajectories), the probability of shifting from a closed habitat to the open (562639 locations), and the probability of shifting from the open to a closed habitat (365684 locations). In all cases, we first tested if open patch availability around individuals significantly explained the variations of the response variable with a likelihood ratio test. If so, it was added in the null model of our model comparison. We then compared what were the best predictors of the variations of the response variable by comparing the null model, the models including the linear variation of average operative temperature in the sun *T_sun_*, and the difference of average operative temperature the sun and the shade *T_diff_*, the models including the quadratic variations of *T_sun_*, and *T_diff_*, and the same models including the interaction of the environmental variable with estimated group size (Table 1). When estimating the probability of shifting from a closed habitat to the open or the probability of shifting from the open to a closed habitat, we fitted the variations of *T_sun_*, *T_diff_*and open patch availability of the previous location (5 minutes earlier) considering that these were more informative of the decision of the bird to shift habitat or not. The best model was identified based on AICc with the “aictab” function from the “AICcmodavg” library (Mazerolle 2025). We accounted for inter-individual and inter-group variations in all response variables by fitting an individual random effect and a group random effect (additively as an individual could change group during the study). Note that we tried to account for temporal autocorrelation between the successive locations of an individual during a day with an AR1 covariance structure when fitting the probability of being in the open. We removed it from models at the end as models did not manage to converge when both temporal autocorrelation and open patch availability were included in the model; our results were robust to the use of one or the other. Probability of being in the open and shifting from one habitat to another were fitted with GLMMs with a binomial error. The same procedures have been run with *T_sun_* and *T_diff_* averaged in the 15 minutes and in the last hour before each location record.

### Variations of distance moved

We calculated the distance moved in one hour by summing the distance between all consecutive locations in that hour to reduce the impact of any noise (e.g. running away from a predator) on the response variable (82324 measurements). The distance moved was square-rooted to approach a Gaussian distribution and fitted with an LMM with a Gaussian error. *T_sun_* and *T_diff_* were averaged per hour, then scaled and centered to facilitate model convergence. Only hour-bouts including 6 or more locations were included and the number of locations in the hour-bout was included in the model, as the fewer the locations the shortest one single displacement may look like (McCann et al. 2021). Model comparison protocol and random effects included in the model were similar as used in the models fitting shifts from one habitat to another.

### Variations of body temperature anomaly

In order to investigate if being active in the sun might impose a cost on thermoregulation, we analyzed the variations of Δ*T_body_* as a function of the quadratic variations of *T_diff_* (as an index of the different temperatures available in the environment) in interaction with the habitat. We accounted for inter-individual variability with a random effect of the individual on the intercept but also for potential effects of the meteorology with a date random effect on the intercept. *T_diff_* was scaled and centered in order to facilitate model convergence.

## Results

Model comparisons results are listed in Appendix 5. The temporal scale of our temperature measurements does not impact our results Appendix 6 and 7.

### Probability of using open habitats

The probability of selecting open patches significantly and strongly increased with open patch availability (χ² = 221286, df = 1, *p* < 0.001; effect: 7.4 ± 0.02). This predicts that if the availability of open patches around the location is very low, the probability of selecting open patches is close to 0, if average it is around 60%, if very high it is almost 100%. We found that the interaction between group size and the instant difference of operative temperatures between sun and shade *T_diff_* best explained the probability to use open habitats (Appendix 5: Table S1). Specifically, the interactions between group size and both the linear term and the quadratic term of *T_diff_* significantly explained the variations in the probability of being in the open (group size*×T_diff_*: χ² = 827.8, df = 2, *p* < 0.001; group size*×T_diff_²*: χ² = 14.3, df = 2, *p* < 0.001). The probability of being in the open was around 60- 65% when temperatures in the sun and in the shade were similar, and decreased with *T_diff_* (Figure 2A). At high *T_diff_*, the probability of being in the open was slightly higher for median groups (*c.* 25%) than for large and small groups (*c.* 15%; Figure 2A).

**Figure 2.**
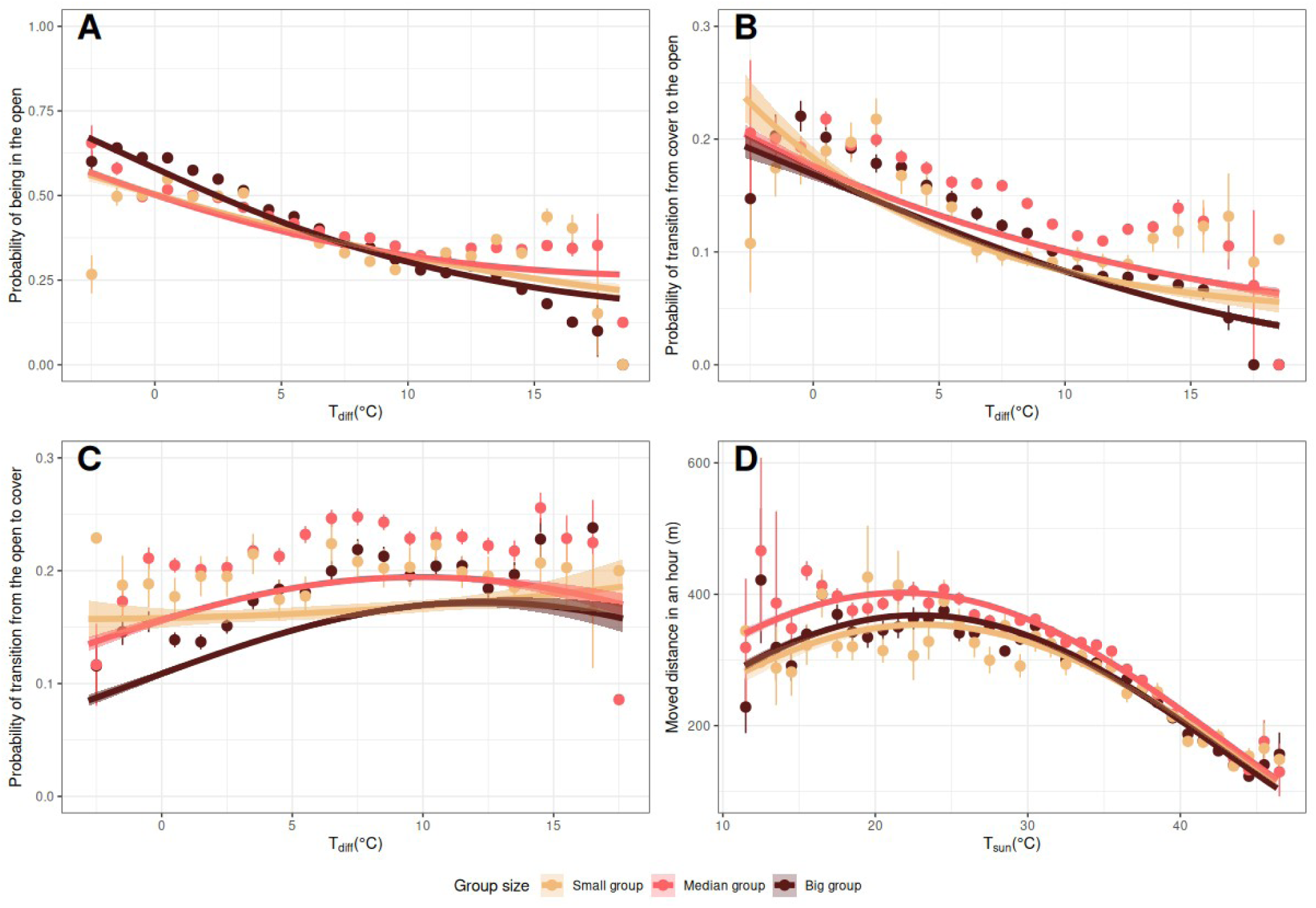
Predictions of the best models identified by model selection for each studied behavior: (A) probability to be in the open vs. cover as a function of *T_diff_*; (B) probability to shift to the open while under cover as a function of *T_diff_*; (C) probability to shift to cover while in the open as a function of *T_diff_* and (D) the distance moved in one hour as a function of *T_sun_*. In all graphs, points are raw proportion of the behavior for each group size for bins of 1°C of *T_diff_* (A,B,C) or *T_sun_* (D). Error bars show the calculated 95% confidence intervals around the calculated proportions. Lines are the predicted probability to do the behavior according to group size. Ribbons are the predicted 95% confidence intervals of this probability. Predictions in A, B and C have been made for an average open habitat availability calculated in the dataset used in the statistical model. The strong predicted effect of open habitat availability explains the deviance of the predictions to the points. In D, predictions have been made for hour-bouts including 12 different locations.

### Probability of moving from cover to open patch

The probability of shifting from a cover to an open patch significantly increased with open patch availability (χ² = 40903, df = 2, *p* < 0.001; effect: 5.1 ± 0.03). The interaction between group size and *T_diff_* best explained the probability to shift to the open when under cover (Appendix 5: Table S2), with the interactions between group size and both the linear term and the quadratic term of *T_diff_* being significant (*T_diff_*: χ² = 106.8, df = 2, *p* < 0.001 ; *T_diff_²*: χ² = 32.4, df = 2, *p* < 0.001). When temperatures in the sun and in the shade were similar, the probability to transition to the open when in the shade was 20-25%, independently of group size (Figure 2B). This probability dropped significantly when *T_diff_* increased. At high *T_diff_*, the probability of shifting to the open was around 6% for small and median groups but was around 3% for large groups. At intermediate *T_diff_*, median sized groups had the highest probability of shifting to open habitats.

### Probability of moving from open to cover patch

The probability of shifting from an open to a cover patch significantly decreased with open patch availability (χ² = 34549, df = 2, *p* < 0.001; effect: -4.5 ± 0.02). The interaction between group size and the quadratic variations of *T_diff_* best explained the probability to shift to a cover habitat from open habitats (Appendix 5: Table S3), with the interactions between group size and both the linear term and the quadratic term of *T_diff_* being significant (*T_diff_*: χ² = 122.5, df = 2, *p* < 0.001 ; *T_diff_²*: χ² = 28.3, df = 2, *p* < 0.001). The probability to transition under cover when in the open increased with *T_diff_* until a threshold of *c. T_diff_*=10°C (Figure 2C). Based on confidence intervals, this probability was significantly higher in median sized groups at intermediate *T_diff_*. Big groups had a lower probability to transition from the open to cover at low *T_diff_*compared to median and small groups (Figure 2C). However, small groups and big groups had similar probability to transition from the open to cover at high *T_diff_* (Figure 2C).

### Distance moved

The interaction between group size and the quadratic variations of *T_sun_* best explained the variations in the hourly distance moved by individuals (Appendix 5: Table S4), with the interactions between group size and both the linear term and the quadratic term of *T_sun_* being significant (*T_sun_*: χ² = 87.2, df = 2, *p* < 0.001; *T_sun_²*: χ² = 7.6, df = 2, *p* = 0.02). The distance moved by an individual in one hour was maximal between 20 and 25°C across all group sizes (Figure 2D). Individuals in median group sizes moved larger distances than other group sizes between 10 and 35°C, whereas distances moved were similar for all group sizes at higher temperatures (Figure 2D). As expected, the number of locations in an hour-bout was positively and significantly correlated with the distance walked in an hour (χ² = 124.3, df = 2, *p* < 0.001; effect: 0.16 ± 0.01).

### Body temperature anomaly

Changes in body temperature (Δ*T_body_*) were significantly predicted by the quadratic value of *T_diff_* in interaction with the habitat (*T_diff_*: χ² = 32.8, df = 1, *p* < 0.001 ; *T_diff_²*: χ² = 13.8, df = 1, *p* < 0.001). Δ*T_body_* was positive when the temperature in the open was higher than the temperature in the shade (*T_diff_* > 0; Figure 3). Δ*T_body_* increased significantly with *T_diff_* and increased more steeply for birds active in the open, resulting in a Δ*T_body_* of c. 0.1°C higher for birds that were active in the open than under cover (Figure 3). At high *T_diff_*, Δ*T_body_* peaked around 0.4°C whatever the habitat where the bird is active (Figure 3).

**Figure 3.**
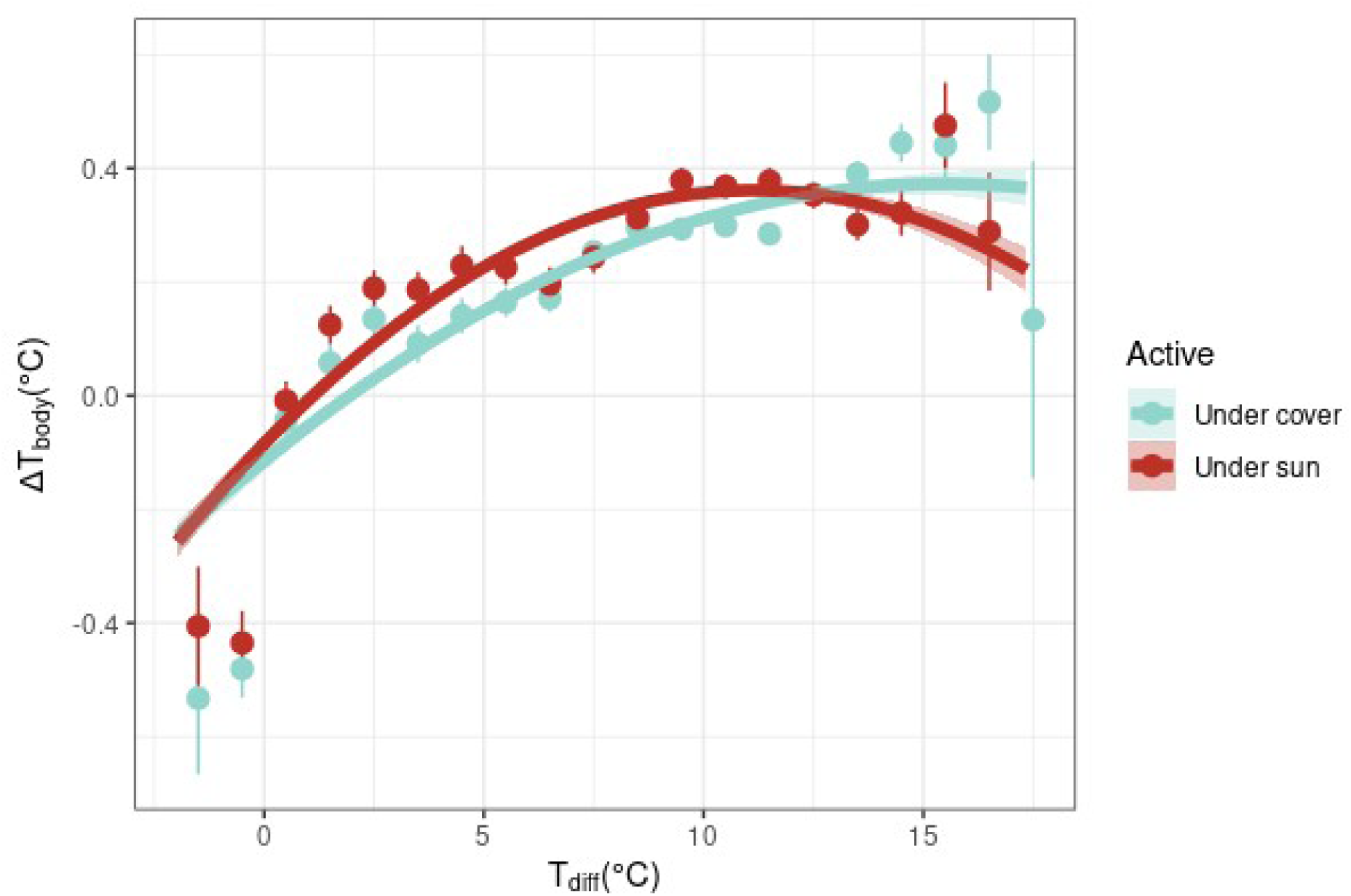
Predicted changes in the deviation between body temperature and resting body temperature (body temperature anomaly Δ*t_body_*) according to *T_diff_* and the habitat (open or cover) where the bird is active. Points are the average Δ*T_body_*for bins of 1°C of *T_diff_* for each habitat. Error bars show the calculated 95% confidence interval around the calculated mean. Lines are the predicted probability to do the behavior according to group size. Ribbons are the predicted 95% confidence intervals of this probability.

## Discussion

We showed that VGFs reduce their use of open habitats—which they rely on for foraging—as temperatures in the sun become higher than temperatures in shade. This effect was captured by a lower probability of presence in the open, a lower probability of transitioning from cover to the open, and higher probability of transitioning from the open to cover under elevated temperatures in the sun. The direction of the effects of thermal constraints on the different habitat use and movement patterns align with a heat avoidance strategy. The use of thermal shelters such as shade to limit overheating is now well-known and expected in larger birds that cannot rely on physiological responses as easily as small passerine birds (Smit et al. 2016; Oswald et al. 2019; Pattinson et al. 2020). Such behaviors mostly involve limiting the exposure of the organism to solar radiations (Wolf and Walsberg 1996, see the operative temperature variations in Figure 1). The main predictor of changes in thermal conditions is the time of the day, with the highest temperatures recorded in the afternoon. Our results mostly suggest that VGFs take advantage of early hours of the day as well as the late ones, but also during cloud cover (*pers. obs.*; see June 5^th^ compared to previous days and the following one in Figure 1; also see for days with more variable weather and more contrasted behavioral responses). In addition, the reduction of the distance moved with increasing temperature indicates that birds spend less time commuting and probably more time resting when foraging patches have high heat exposure, which is consistent with a strategy aiming at avoiding both hot micro-habitats but also when lowering metabolic rate and thus heat production (Clark 1987; Pattinson et al. 2020). We also observed VGFs panting (or doing other heat dissipation behavior such as wing dropping or fluttering) while resting under shade in the hottest hours of the day, as expressed by many other arid habitats birds (Pattinson et al. 2020). This suggests that VGFs compensate foraging activity under heat by more resting and heat dissipation behaviors, as a result of the trade-off between foraging and heat dissipation. However, the lower distance moved observed at lower temperatures (when birds were mostly in the open) also suggests that birds move less when actively foraging in low heat conditions (i.e. because they remain on the relatively small glades). These observations together suggest that physiological responses (especially evaporative water losses, EWL) alone are insufficient for the regulation of body temperature under the high and frequent heat conditions of the dry season or that high limitations in water availability during the dry season impose a higher cost on evaporative water losses and shift bird thermoregulation processes from physiology to behavior (Pattinson et al. 2020).

Our results also show that active birds have an increase of their body temperature of c. 0.3°C compared to period of inactivity, as a result of the trade-off between heat dissipation and activity. This predicted hyperthermia is, however, small compared to the variations observed in body temperature across the day or amongst individuals, which can range to several degrees (Figure S5). The 0.1°C difference of body temperatures between bouts of activity in the open or under cover also confirm the impact of radiation in the heat constraints experienced by individuals (Wolf and Walsberg 1996). Overall, the range of body temperatures observed in our study do not exceed expected body temperature of such a large bird (McKechnie and Wolf 2019) as well as it is coherent with the range of temperatures observed in a closely related species (Withers and Crowe 1980) and far from the range of temperatures observed under both pathological or facultative hyperthermia (Gerson et al. 2019). The behavioral modulation of heat constraints is thus likely to be efficient at mitigating the costs of heat on the organism. Heat management in VGFs is also likely to be supported by other strategies, such as panting, and morphological adaptations, such as bare heads allowing heat dissipation despite their dark plumage and reduces the costs of foraging under heat (Galván, Palacios, and Negro 2017). However, the consequences of these behaviors in water losses and thus dehydration—and in turn individual fitness—remains a blind spot of our study (Albright et al. 2017; Rozen-Rechels et al. 2019; Czenze et al. 2020).

We observed that the changes in birds’ behaviors were also modulated by their group size. We found consistent group size effects, with intermediate (i.e. *optimal*; Bertram 1978) group sizes often expressing more flexible responses (e.g. higher transition probabilities to and from cover; Sibly 1983; Papageorgiou and Farine 2020b). Large groups seemed more inclined to avoid open habitats at high heat constraints while intermediate-size groups (median group sizes) seemed to have more opportunities to move and shift habitats when heat increased, and more particularly at intermediate temperatures. This is consistent with recent observations that intermediate-size groups of the same population are the most mobile (Papageorgiou and Farine 2020b). This is likely due to the time costs and coordination challenged associated with making decisions by larger groups (Davis, Crofoot, and Farine 2022). In other words, when faced with heat constraints in the sun, large groups are likely to be less able to respond quickly by shifting between habitat conditions. By contrast, small groups show similar habitat use patterns as intermediate-size groups under high heat constraints, suggesting that they can benefit from making faster decisions in response to dynamic changes in their (thermal) environment. Overall, we therefore expect that an increase in the length and frequency of hot dry seasons in such savanna habitats could favor smaller groups despite the benefits against predation in large groups. Alternatively, we could also interpret the tendency by larger groups to avoid open patches at higher temperatures as an ability to take a more optimal decision in terms of thermoregulation and avoid warmer habitats more efficiently.

## Conclusion

Beyond the result that vulturine guineafowl habitat-use behavior is driven by heat avoidance, which might therefore constrain foraging behaviors, we have shown that the behavioral responses of individuals to heat are complex. One particularly important dimension of our findings is that thermoregulatory behaviours—or strategies—appear to be related to group size. At the relatively short time scale of our study, we found that VGFs cope with heat efficiently, however we lack information on the fitness consequences of these behavioral modulations on individual survival and recruitment (but note that the subsequent drought that continued until 2023 resulted in total suppression of reproductive behaviors due to a lack of dense cover for nesting or the lack of food resources). Thus, our results underline the importance of sociality on individual-level responses to unfavorable weather conditions, a factor that is often overlooked in studies of the responses of individual organisms to heat. A better understanding of the consequences of climate changes on animals, and more generally on biodiversity, will also benefit from greater consideration of the social landscape (Webber et al. 2023).

## Supporting information

All appendices

## Acknowledgments

All procedures were approved by the Ethics Council of the Max Planck Society (2016_13/1), the National Commission for Science, Technology and Innovation (annual permits for all contributors of this study who worked in Kenya, licence NACOSTI/P/21/7442 for David Rozen-Rechels in 2021), Kenya Wildlife Service (annual permits for research and capture, KWS/904), the National Environment Management Authority (NEMA/AGR/68/2017), the Wildlife Research and Training Institute (WRTI-0026-02-21), and the National Museums of Kenya (NMK/ZLG/TRN/6.5). The procedure of implantations of ECG loggers was reviewed by the Animal Welfare Officer at the University of Zurich. The procedure was realized by Dr. Daniel Zuñiga and performed under the supervision of Dr. Maureen Kamau from the Kenyan Wildlife Service. The procedure was reviewed and Dr. Daniel Zuñiga was approved to perform the surgeries by the Kenya Veterinary Board (KVB/FVS/Voll/6).

We thank Alex Baiywa, Wismer Cherono, Janet Wangare Kariuki, Mary Waithira Ngugi, Edel Awour Odhiambo, Monicah Wambui, John Wanjala, and all the field assistants and all the rangers, administrative people in Mpala Research Center allowing this research to be conducted since 2016. This research was funded by the European Research Council (ERC) under the European Union’s Horizon 2020 research and innovation programme (grant agreement No. 850859 awarded to DRF), the Swedish Research Council (Grant Number 2019-06407 awarded to CHW), and an Eccellenza Professorship Grant of the Swiss National Science Foundation (Grant Number PCEFP3_187058 awarded to DRF). DRR received additional funding from a Humboldt Postdoctoral Fellowship granted to David Rozen-Rechels.

## Authors contributions

DRR elaborated the hypotheses, set-up the thermal study, implemented the remote sensing mapping of the landscape and the temperatures measurements, collected behavioral data, analyzed the data and wrote the manuscript. BN advised the implementation of the protocols and managed the collection of GPS, census and operative temperature measurements. DP gave expertise on the hypotheses and the implementation of the study as well as analyses. MO managed the determination of communities. NB analyzed all videos. CHW implemented the ECG loggers study. JKI helped calculating group size. PN supervised the long-term monitoring of guineafowl. DRF established the guineafowl project, led the project, and supervised the study. All authors contributed to the writing of the manuscript.

