## Supplementary material for "Group size influences behavioral plasticity in responses to thermoregulation-foraging trade-offs by a socially cohesive bird": All appendices

**Appendix 1 – Differences of foraging effort between open and cover patches**

All statistical tests described below have been made in R (4.2.1 “Funny-Looking Kid”). Generalized linear mixed models (GLMM) were fitted with the “glmmTMB” library, residuals distribution and homoscedasticity as well as the leverage effects of outliers or overdispersion of data were assessed with the “DHARMa” library. All model were simplified using loglikelihood ratio tests comparing models using a Maximum Likelihood (ML) estimation. Final models were analyzed after being fitted with a Random Effect Maximum Likelihood (REML) estimation.

*Open (vs. cover) patch use around glades and foraging propensity*

From December 7^th^ 2020 to February 13^th^ 2021, one observer scan-sampled the open vs. cover patch selection and the behaviors around glades of groups of guineafowl in 15 different locations (the same group could have been sampled twice, but many times different groups were mixed up or we only observed a subset of a group, as a consequence we ignored any group effect in the following analyses). From 06:00 to 09:00, from 09:00 to 12:00, from 12:00 to 15:00 or from 15:00 to 17:30 we looked after groups of birds in different parts of the landscape of study. As soon as a group was found, we stayed with the group as long as possible (we stopped recording behaviors when the group left and we were not able to follow it in the bush or when we reached the end of the session). Every 5 minutes, the observer counted the number of birds visible and among them the number of birds in the open in the sun, the number of birds in the open in the shade, and the number of birds under cover of vegetation. Within each micro-habitat category we counted the number of birds foraging (pecking, scratching the ground with their head, moving slowly around looking on the ground, sliding their beak on a grass) versus birds having any other behavior (i.e. resting, having agonistic behavior with other birds, preening, being vigilant...). The observer also noted if the weather was sunny or cloudy at the same time. When cloudy, birds in the open were all counted as in the shade. If the count was disturbed by an uncontrolled event (something made the birds ran away, the group decided to leave the area and started to be on the move), we discarded the scans as long as the behavior of the birds were disturbed.

We analyzed how open vs. cover patch use (here measured as the number of birds in a open patch vs. in a cover patch) changed according to a quadratic fit of the time of the day as well as the additive effect of the total number of birds counted and the weather (sunny vs. cloudy) with a GLMM with a binomial error and a random effect of the date on the intercept to fit the variance explained by differences between days. Open (vs. cover) patch use variation was significantly explained by a quadratic fit of the time of the day (time: χ² = 16.4, df = 1, p < 0.001; time²: χ² = 67.4, df = 1, p < 0.001). The use of open patch was maximal (and close to one) early in the morning and late in the day, and minimal around noon (Appendix 1: Figure S1). Birds were also more likely to be found in the open when numerous (χ² = 71.1, df = 1, p < 0.001, estimate: 0.05 ± 0.006) but less likely when sunny (χ² = 12.2, df = 1, p < 0.001, estimate: -0.67 ± 0.19).

We explained the variance in the propensity to forage (here measured as the number of birds showing foraging birds vs. other behaviors) with the interaction between the patch type (open vs. cover) in interaction with the quadratic variations of time and an additive effect of the weather (sunny vs. cloudy) with a GLMM with a binomial error and a random effect of the date on the intercept to fit the variance explained by differences between days. Overdispersion and deviation of residuals to a Gaussian distribution were corrected using an observation level random effect. The propensity to forage is significantly explained by the interaction between the quadratic term of the variation of the time of the day and the patch type (χ² = 4.1, df = 1, p < 0.001). Foraging propensity is high all along the day in the open (around 80%) while it drops to less than 50% under cover around midday (Appendix 1: Figure S2). The weather did not explain variations in foraging propensity (χ² = 0.01, df = 1, p = 0.92).

*Pecking rate in the open and under cover*

At the same time, between each scan, we recorded videos of individual birds in both open and cover patches. We recorded 55 individuals at least twice (at least once in an open patch and once under cover) in videos lasting in average 94s (median: 114s, minimum: 13s, maximum: 201s). One observer analyzed all videos. First, all bouts of the videos in which the behavior could not be identified (when hidden by the vegetation or when the orientation of the birds did not allow to see its head) were excluded resulting in analyzable videos lasting in average 86s (median: 96s, minimum: 10s, maximum: 201s). Second, the observer identified bouts of videos when the bird was foraging, i.e. looking on the ground or in the vegetation for food, pecking, scratching the ground. In this part were excluded bouts corresponding to commuting (neck straight up for more than 2 seconds), interactions (running to other birds) or bouts when the bird was resting (doing nothing, preening, scratching its body and/or laying down). Birds were foraging from 2% to 100% of the duration of the videos. Third, the observer counted the number of pecks, used here as a proxy of foraging effort, during the foraging bouts using the Tally Count app. When pecking happened to be fast, video speed was slowed down to 50%. In case of uncertainty in the counts, the counting was repeated several times and the average number of counts was used in further analyses (if discrepancy was high, the number used was the one with the best consensus).

We then analyzed how peck rate changed along time (hour of the day) according to the patch: cover or open. We calculated peck rate for each video as the ratio of pecks count and foraging bouts duration. We square-rooted peck rates in order that their distribution approached a gaussian distribution. We analyzed the variations of square-rooted peck rates with a GLMM fitting the quadratic and linear terms of time, both in interaction with the patch type. The individual identity and the date were fitted as random effects on the intercept in order to account for interindividual differences in peck rates and potential differences of food availability or behavior explained by the session.

Root-squared peck rate variations were significantly explained by the interaction between the patch type and the quadratic term of the time (χ² = 7.00, df = 1, p = 0.008). Peck rate was higher in the open (ca 1.1 peck/s) than under cover (ca 0.6 pecks/s) except at the end of the day when it got approximately similar in both patch type (Appendix 1: Figure S3). Note that the statistical model showed a significant deviation of the distribution of residuals from a gaussian distribution but the results remained robust when fitting peck rate calculated with the total video duration with a model showing no issues with residuals.

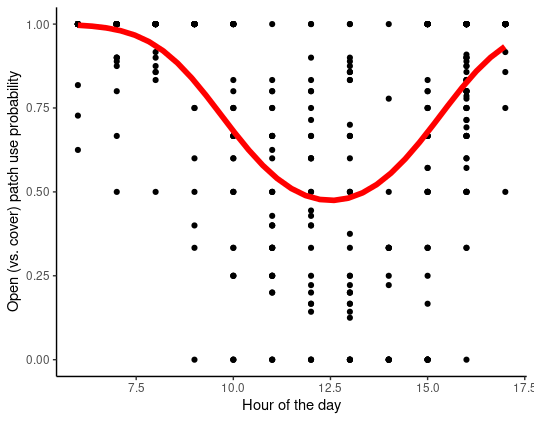

Appendix 1: Figure S1: Changes in open (vs. cover) patch use probability over the course of the day. Black dots show calculated proportion of birds using open (vs. cover) patch at each counts. The red line represents the predicted probability of open (vs. cover) patch use according to the hour of the day from the best model for the average number of birds counted and a sunny day.

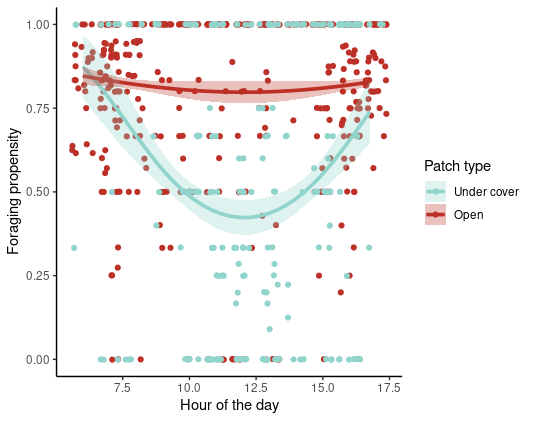
Appendix 1: Figure S2: Foraging propensity according to the hour of the day and the patch type (red: open, blue: cover). Dots represent calculated proportion of birds foraging at each scan. Lines are predicted foraging propensity according to the hour of the day and the patch type from the best model, ribbons represent 95% confidence intervals.

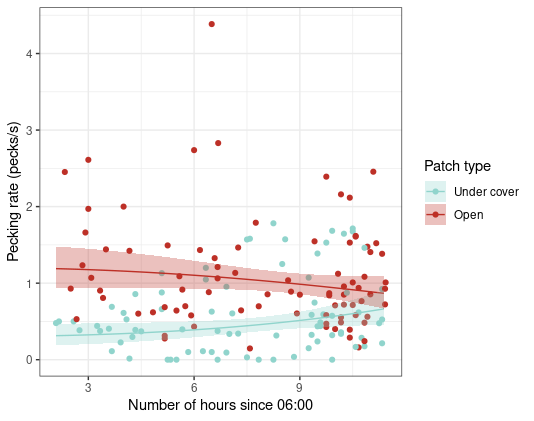
Appendix 1: Figure S3: Pecking rate (pecks/s) according to the hour of the day and the patch type (red: open, blue: cover). Dots represent raw peck rates for each video. Lines are predicted peck rate according to the hour of the day and the patch type from the best model, ribbons represent 95% confidence intervals.

**Appendix 2 – Remote sensing and landscape description**

In this project, we aim at calculating diverse indices of vegetation presence based on remote sensing data in order to able to interpolate open (grassland or naked soil) patches and covered (trees and bush) patches. Sentinel-2 mission consists of 2 satellites (A and B) continuously recording multi-spectral pictures, with a 10 days frequency and a 5 days difference between them. Each area pictured, of an area of10000 km² is called a tile. We focus on tile 37N BA containing Mpala Research Center (MRC). Sentinel 2 data can be downloaded in two different format: L1C and L2A. The main difference between them is the atmospheric correction which is already processed for L2A. However, L2A data are only available starting in 2019. As we needed remote sensing data from the onset of our study in MRC (2017) to now for other projects, we decided to download L1C data. All 37N BA tile photographs have been downloaded in the Google Storage of Sentinel-2 data with the *gsutil* program (https://cloud.google.com/storage/docs/public-datasets/sentinel-2?hl=en

) from the earliest date possible (October 11^th^ 2017) to the end of this project (August 21^st^ 2021). We then used Sen2Cor standalone 2.8 for atmospheric corrections with default parameters in order to produce L2A data for all downloaded tiles.

We sampled and subset the L2A tiles with the *ESA SNAP* program in command line. We resampled all tiles at the 10m resolution (some bands have lower resolution). We only kept the band of interest for future actions: Band 3 (green), Band 4 (red), Band 8 and 8A (near infrared NIR, 8A is more restrictive and is the one used later), Band 11 (Short-Wave IR SWIR), and the “quality scene classification” band. We also cropped all tiles in order to optimize computing operations (minimum longitude: 36.84°E, maximum longitude: 36.98°E, minimum latitude: 0.24°N, maximum latitude: 0.36°N; the coordinates used are large enough to contain the whole study area). The code used can be found on the Zenodo repository (url to be provided upon publication). For each date/tile we obtained a stack of rasters.

Unless otherwise stated, the procedures below have all been performed with R version 4 software and the raster, rgdal, and sf libraries. We then cropped all obtained stacks of rasters using a polygon mask containing all locations of the birds included in the study while removing some part of the landscape in altitude in which birds only commute and where the soil is of different composition, making habitat identification less reliable (Appendix 2: Figure S1). Finally, we identified all pixels that have to be excluded for vegetation indices calculations based on the “quality scene classification” band: dark features, cloud shadows, unclassified, cloud medium probability, cloud high probability, thin cirrus, snow or ice. If these pixels covered more than 20% of the map, the entire map was excluded from the study. In the other case, we only excluded the concerned pixels from the analysis. For each stack of rasters, i.e. each date, we then calculated NDVI raster maps (Normalized Difference Vegetation Index, eqn. 1, Appendix 2: Figure S2), NDWI maps (Normalized Difference Water Index, water content in vegetation, eqn. 2, Gao 1996) and brightness maps (eqn. 3, Valero et al. 2016).

Eqn. 1 $NDVI=\frac{NIR-Red}{NIR+Red}$

Eqn. 2 $NDWI=\frac{NIR-SWIR}{NIR+SWIR}$

Eqn. 3 $Brightness=\sqrt{{Green}^{2}+{Red}^{2}+{NIR}^{2}+{SWIR}^{2}}$

with NIR meaning Near InfraRed, SWIR meaning Short-Wave InfraRed. We summarized the indices by calculating statistics maps such as average, standard deviation, maximum, minimum and median NDVI, NDWI and brightness maps.

We then used real colors satellite pictures of the landscape in order to create a training set of polygons identified as being located in open (grass or naked soil under sun when there is no cloud cover) or cover (under bush or trees, mostly under shade whatever the weather) patches. We draw by hand 80 polygons of each type, as couple (one open patch always close to a cover patch), in order to sample the integrity of the landscape of the study. This procedure has been realized in QGIS 3.

All pixels of the NDVI, NDWI, and brightness statistics maps included in these polygons have been then extracted and identified as open (823 pixels) or cover (1034 pixels). We could then relate the NDVI, NDWI and brightness statistics to the habitats. To do so, we ran a Random Forest procedure in which these pixels have been used as training sets (Breiman 2001). This procedure generates decision trees based on a subset of a training set of samples. A decision tree split these objects based on their variables (such as average NDVI or maximum brightness) in order to classify them in different categories. At each node of a tree, a variable is selected as discriminant and a threshold is defined as a rule of decision. The final nodes (or leaf nodes) are the class aimed for identification (open or cover in this case). The Random Forest procedure select the decision tree that occurred the most frequently in the different subsets. It also provides an error estimate (Our of Bag OOB Error) which is the proportion of the samples out of the subset of samples used to generate the decision tree (out of bag) which are not identified in the right class based on the tree. It also provides the relative importance of each variables with two indices: the Mean Decrease Accuracy (difference of correctly identified sample between original trees and tress with variables decision thresholds permuted) and Mean Decrease Gini (Gini index measures how “pure” a node is, i.e. it allows the discrimination of one class; Mean Decrease Gini measures the sum of Gini decrease for all nodes for one variable over all trees). The procedure requires that the number of occurrence of each habitat is equilibrated. We then randomly picked 823 cover pixels within the 1034 pixels from the training sample. We tuned a Random Forest procedure using the randomForest R library with 250 tested decision trees. The OOB error of the decision tree selected is 2.05%. 1.7% of the cover pixels have been misidentified as open and 2.4% of open pixels have been misidentified as cover. Mean Decrease Accuracy and Mean Decrease Gini showed that average and median brightness, as well as median NDVI were the most discriminant variables (Appendix 2: Figure S3).

In order to validate the classification, we randomly generated 200 points in the landscape, 100 in each class of habitat (open or cover). Using Google Earth, we blindly labeled each points as cover, open (or undetermined as the picture resolution was sometimes not fine enough to visually identify the habitat). The accuracy of habitat class prediction by Random Forest was then estimated at 82%. All known glades in the landscape were correctly identified. We also predicted the habitat class using another method called Support Vector Machine (Noble 2006) which led to the same classification accuracy and was 80% similar to Random Forest predictions (see codes on Zenodo for details). We used the habitat class map predicted by Random Forest in the rest of the study (Appendix 2: Figure S4). In addition, we calculated the raster of the proportion of open pixels in a 240m square centered on each focal pixel (99%-quantile of the distribution of the distance moved by birds in 5min is *c.* 120m), herafter called “open patch availability” (Appendix 2: Figure S5).

Breiman, Leo. 2001. “Random Forests.” *Machine Learning* 45 (1): 5–32. <https://doi.org/10.1023/A:1010933404324>.

Gao, Bo-cai. 1996. « NDWI—A normalized difference water index for remote sensing of vegetation liquid water from space ». *Remote Sensing of Environment* 58 (3): 257‑66. <https://doi.org/10.1016/S0034-4257(96)00067-3>.

Noble, William S. 2006. “What Is a Support Vector Machine?” *Nature Biotechnology* 24 (12): 1565–67. <https://doi.org/10.1038/nbt1206-1565>.

Valero, Silvia, David Morin, Jordi Inglada, Guadalupe Sepulcre, Marcela Arias, Olivier Hagolle, Gérard Dedieu, Sophie Bontemps, Pierre Defourny, and Benjamin Koetz. 2016. “Production of a Dynamic Cropland Mask by Processing Remote Sensing Image Series at High Temporal and Spatial Resolutions.” *Remote Sensing* 8 (1): 55. <https://doi.org/10.3390/rs8010055>.

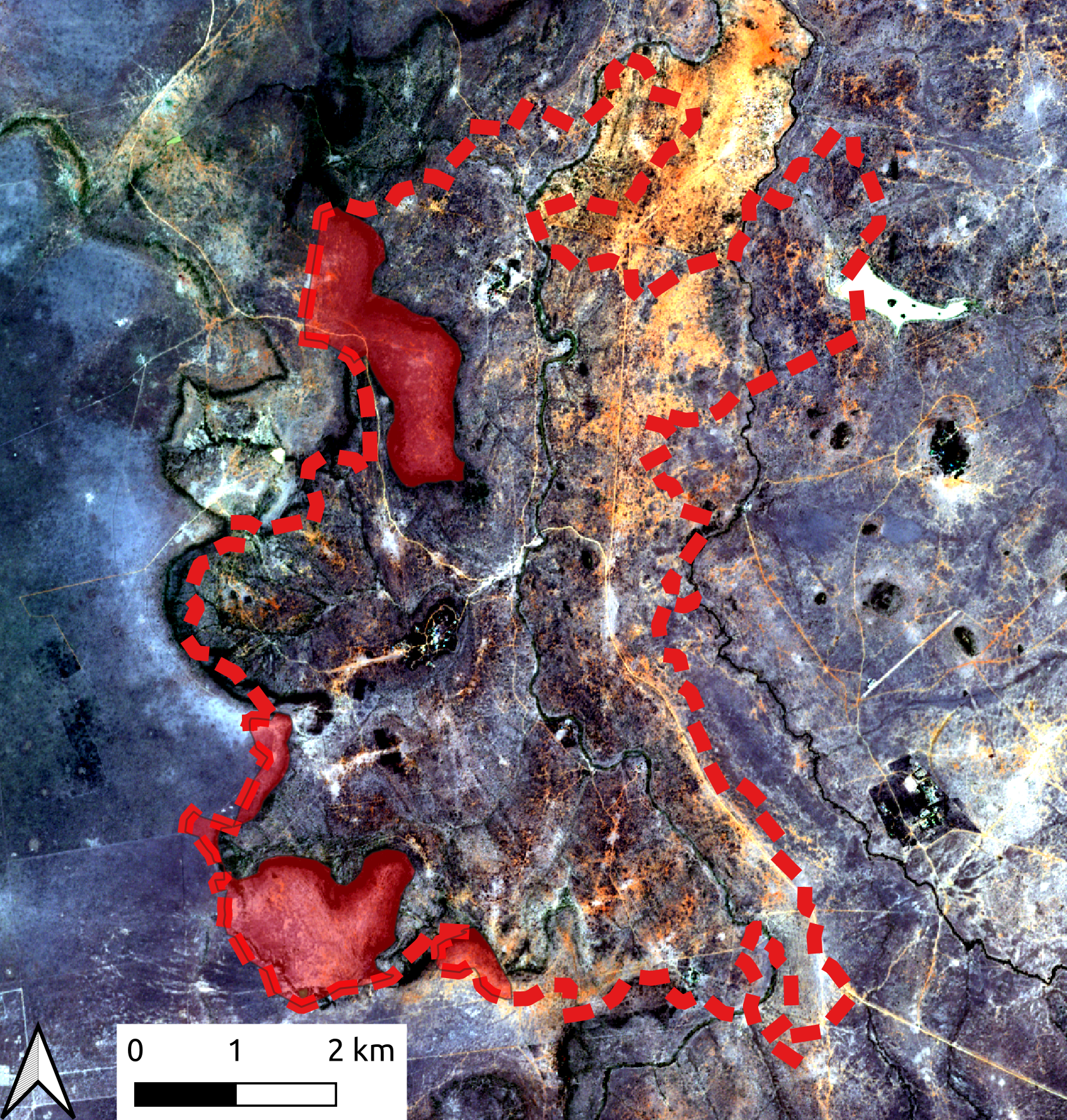
Appendix 2: Figure S1: Sentinel-2 photographs (RGB) of the studied landscape define as the zone containing all the locations of the birds during the study (dashed red). Plateaus having a very different vegetation, Random Forest procedures are less efficient in predicting open and cover habitats and these areas (transparent red) have been consequently excluded from the analyses.

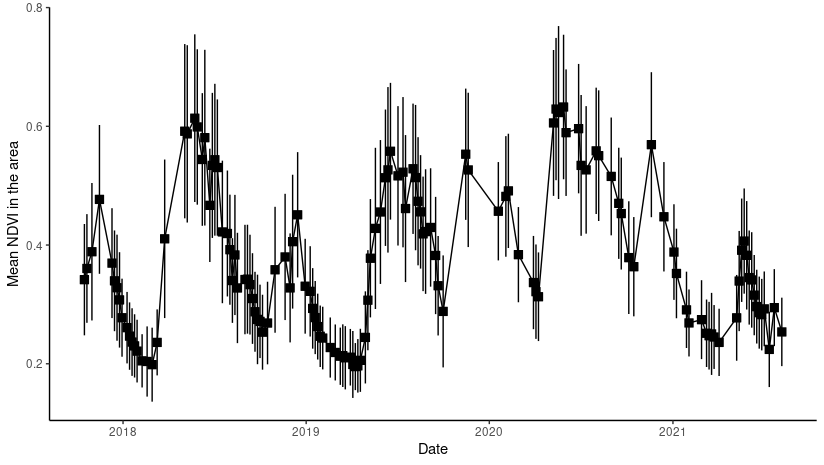
Appendix 2: Figure S2: Variations of NDVI in the studied landscape (Appendix 2: Figure S1). Our period of focus (from February to July 2021) is marked by a particularly long drought that had not end by the end of the study.

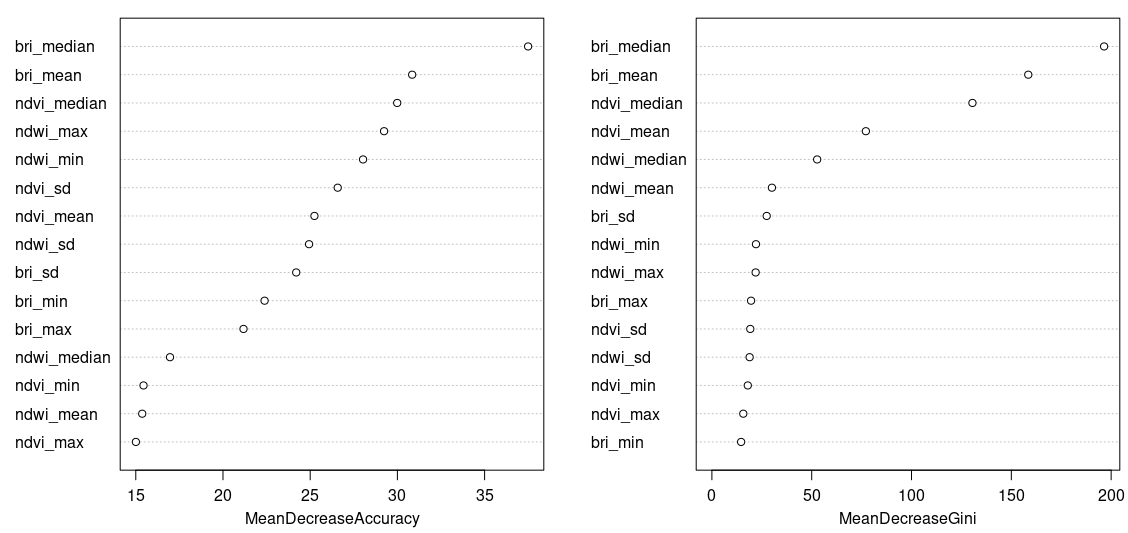
Appendix 2: Figure S3: Mean Decrease Accuracy and Mean Decrease Gini for all variables extracted from the Random Forest procedure. bri = Brightness, sd = Standard Deviation, min = minimum, max = maximum.

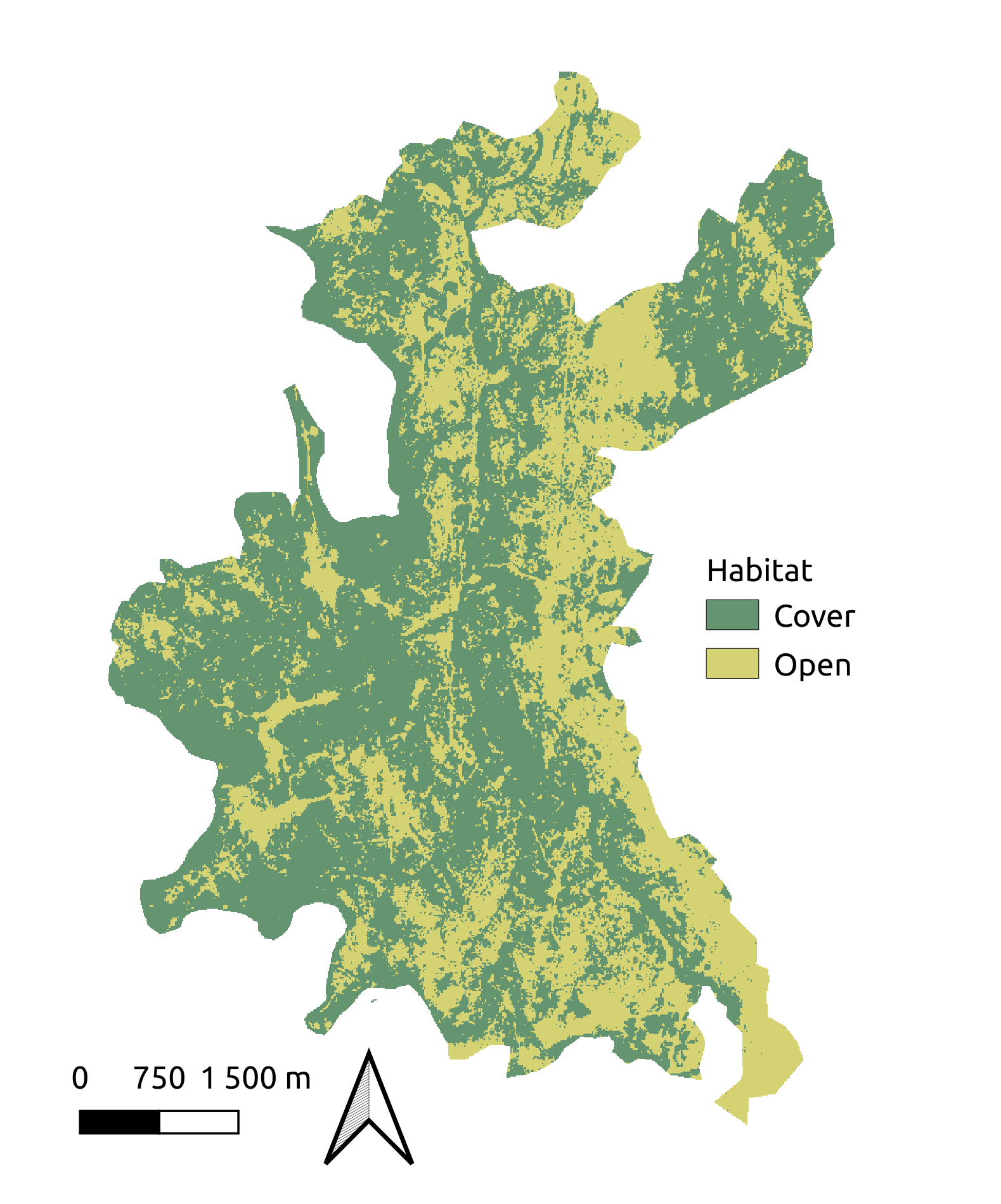
Appendix 2: Figure S4: Prediction of open and cover patches with Random Forest Procedure.

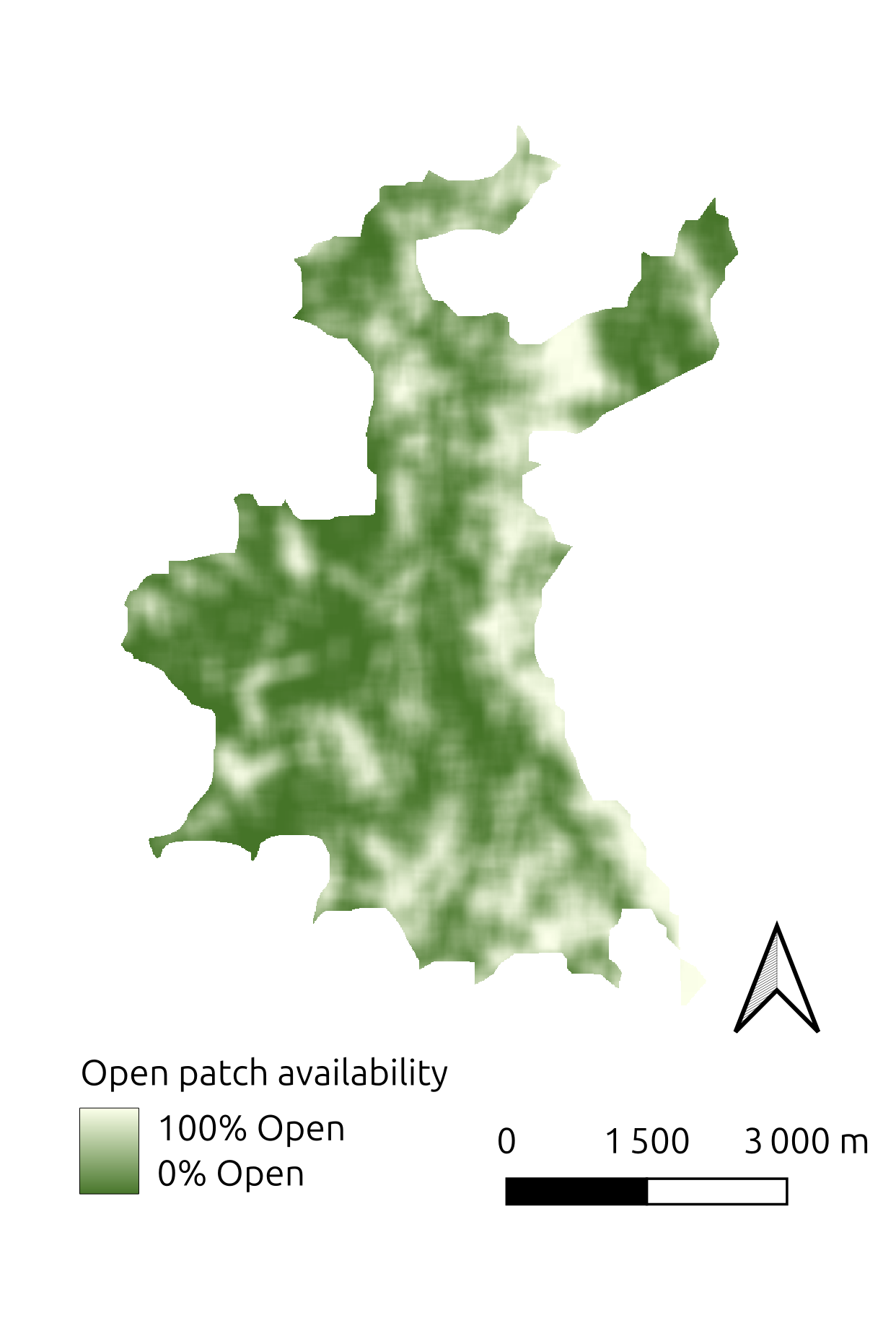
Appendix 2: Figure S5: Proportion of open pixels (“open patch availability”) from Figure SX in 240m-square centered on each pixel (i.e. in c. 120m radius buffer centered on each pixel). The contours of the map are cropped on a 120m distance as focal pixel calculations including pixels out of the studied landscape (Figure SX and SX) were excluded.

**Appendix 3 – Supplementary Information on body temperatures**

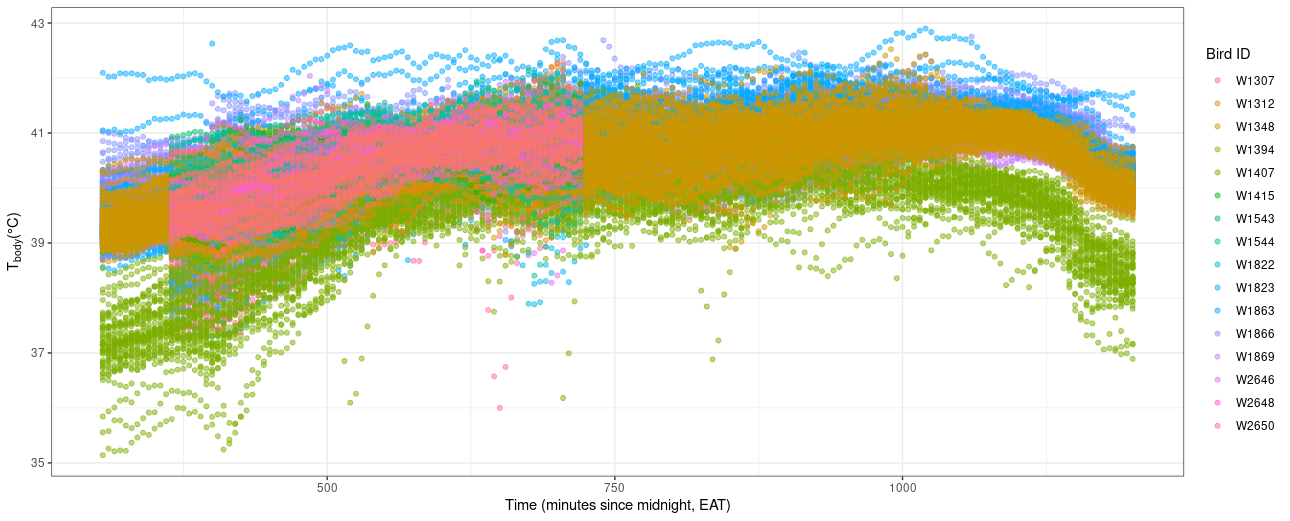
Appendix 3: Figure S1: Body temperatures T_body_ averaged every 5 minutes for all 16 birds. Colors represent birds identity.

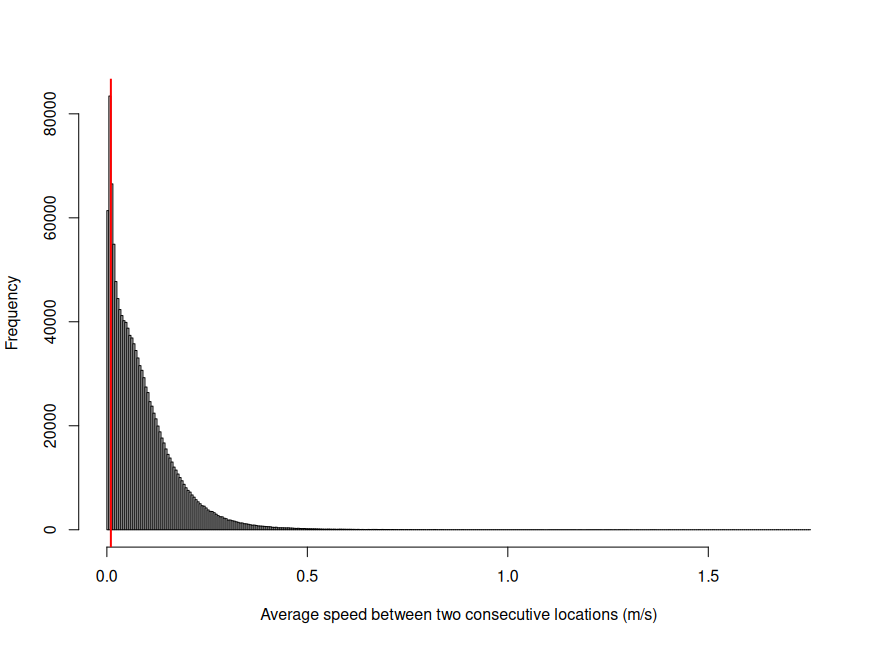
Appendix 3: Figure S2: Distribution of the average speed between two consecutive locations in m/s.

**Appendix 4 – Operative temperature variations**

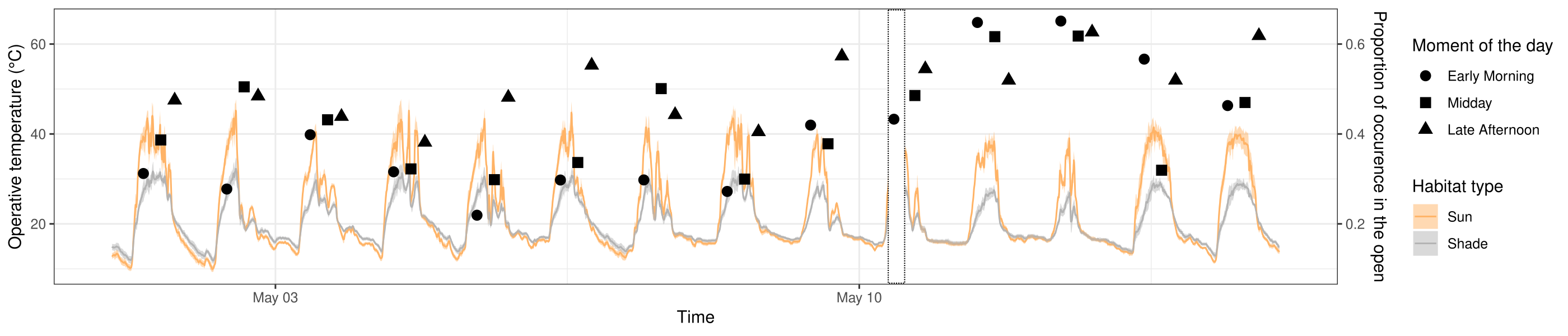
Appendix 4: Figure S1: Variations of the operative temperature in the open averaged for all models in the open (i.e. *T_sun_*) and operative temperature in cover averaged for all models in cover, for a subset of dates (for clarity). Ribbons show the standard deviation. Points are showing the proportion of occurrence in the open of VGFs for 3 separated moment of the day indicated by shape: early in the morning (circles, 07:00 to 08:59 EAT), midday (squares, 12:00 to 13:59 EAT), and late afternoon (triangles, 16:00 to 17:59 EAT). The white rectangle with the black dotted line border shows data removed during the process of replacement of the ibuttons (once every two weeks).

**Appendix 5 – Model comparison results**

| Model # | Explicative variables | K | AICc | ΔAICc | Loglikelihood |
| --- | --- | --- | --- | --- | --- |
| 10 | [*T_diff_* + *T_diff_*²] *× Gs* | 12 | 891649 | 0 | -445813 |
| 9 | *T_diff_ × Gs* | 9 | 891768 | 119 | -445875 |
| 5 | *T_diff_* + *T_diff_*² | 6 | 892682 | 1033 | -446335 |
| 4 | *T_diff_* | 5 | 892807 | 1158 | -446399 |
| 8 | [*T_sun_* + *T_sun_*²] *× Gs* | 12 | 894480 | 2831 | -447228 |
| 7 | *T_sun_ × Gs* | 9 | 895423 | 3774 | -447703 |
| 3 | *T_sun_* + *T_sun_*² | 6 | 895519 | 3870 | -447753 |
| 2 | *T_sun_* | 5 | 896422 | 4773 | -448206 |
| 6 | *Gs* | 6 | 913754 | 22104 | -456871 |
| 1 | 1 | 4 | 913828 | 22178 | -456910 |

Appendix 5: Table S1: Model comparison based on AICc of the model fitting the variations of the probability of being in the open. The models were fitted with the glmmTMB library, and included open patch availability as fixed effect in order to account for habitat availability around each location, as well as an individual and a group random effect on the intercept. The model number refers to the hypotheses and models listed in Table 1 of the main manuscript. K stands for the number of fitted parameters.

| Model # | Explicative variables | K | AICc | ΔAICc | Loglikelihood |
| --- | --- | --- | --- | --- | --- |
| 10 | [*T_diff_* + *T_diff_*²] *× Gs* | 12 | 379421 | 0 | -189698 |
| 9 | *T_diff_ × Gs* | 9 | 379470 | 50 | -189726 |
| 5 | *T_diff_* + *T_diff_*² | 6 | 379556 | 135 | -189772 |
| 4 | *T_diff_* | 5 | 379577 | 157 | -189784 |
| 8 | [*T_sun_* + *T_sun_*²] *× Gs* | 12 | 380026 | 606 | -190001 |
| 3 | *T_sun_ + T_sun_²* | 6 | 380099 | 678 | -190043 |
| 7 | *T_sun_ × Gs* | 9 | 380864 | 1443 | -1900423 |
| 2 | *T_sun_* | 5 | 380921 | 1501 | -190456 |
| 6 | *Gs* | 6 | 385349 | 5929 | -192669 |
| 1 | 1 | 4 | 385358 | 5938 | -192675 |

Appendix 5: Table S2: Model comparison based on AICc of the model fitting the variations of the probability of transition to the open when under cover. The models were fitted with the glmmTMB library, open patch availability at the previous location as fixed effect in order to account for habitat availability around each location,and included an individual and a group random effect on the intercept. The model number refers to the hypotheses and models listed in Table 1 of the main manuscript. K stands for the number of fitted parameters.

| Model # | Explicative variables | K | AICc | ΔAICc | Loglikelihood |
| --- | --- | --- | --- | --- | --- |
| 10 | [*T_diff_* + *T_diff_*²] *× Gs* | 12 | 320431 | 0 | -160204 |
| 5 | *T_diff_* + *T_diff_*² | 6 | 320573 | 142 | -160281 |
| 9 | *T_diff_ × Gs* | 9 | 320581 | 149 | -160281 |
| 4 | *T_diff_* | 5 | 320714 | 283 | -160352 |
| 8 | [*T_sun_* + *T_sun_*²] *× Gs* | 12 | 320992 | 561 | -160484 |
| 7 | *T_sun_ × Gs* | 9 | 321040 | 609 | -160511 |
| 3 | *T_sun_* + *T_sun_*² | 6 | 321134 | 703 | -160561 |
| 2 | *T_sun_* | 5 | 321167 | 735 | -160578 |
| 6 | *Gs* | 6 | 321473 | 1041 | -160730 |
| 1 | *1* | 4 | 321475 | 1043 | -160733 |

Appendix 5: Table S3: Model comparison based on AICc of the model fitting the variations of the probability of transition under cover when in the open. The models were fitted with the glmmTMB library, open patch availability at the previous location as fixed effect in order to account for habitat availability around each location, and included an individual and a group random effect on the intercept. The model number refers to the hypotheses and models listed in Table 1 of the main manuscript. K stands for the number of fitted parameters.

| Model # | Explicative variables | K | AICc | ΔAICc | Loglikelihood |
| --- | --- | --- | --- | --- | --- |
| 8 | [*T_sun_* + *T_sun_*²] *× Gs* | 13 | 460457 | 0 | -230216 |
| 3 | *T_sun_* + *T_sun_*² | 7 | 460616 | 159 | -230301 |
| 10 | [*T_diff_* + *T_diff_*²] *× Gs* | 10 | 463960 | 3503 | -237970 |
| 7 | *T_sun_ × Gs* | 6 | 464084 | 3627 | -232036 |
| 5 | *T_diff_ + T_diff_²* | 13 | 464614 | 4157 | -232294 |
| 9 | *T_diff_ × Gs* | 10 | 464736 | 4279 | -232358 |
| 2 | *T_sun_* | 7 | 464755 | 4298 | -232371 |
| 3 | *T_diff_* | 6 | 464867 | 4410 | -232428 |
| 6 | *Gs* | 7 | 478773 | 18316 | -239379 |
| 1 | 1 | 5 | 478813 | 18356 | -239401 |

Appendix 5: Table S4: Model comparison based on AICc of the model fitting the variations of the distance moved in one hour. The models were fitted with the glmmTMB library, and included the number of locations used to calculate the distance in the hour bout, an individual and a group random effect on the intercept. The model number refers to the hypotheses and models listed in Table 1 of the main manuscript. K stands for the number of fitted parameters.

**Appendix 6 – Robustness of results of replacing instant temperature measurements to the average temperature conditions in the last 15min**

| Model # | Explicative variables | K | AICc | ΔAICc | Loglikelihood |
| --- | --- | --- | --- | --- | --- |
| 10 | [*T_diff_* + *T_diff_*²] *× Gs* | 12 | 892121 | 0 | -446048 |
| 9 | *T_diff_ × Gs* | 9 | 892172 | 52 | -446077 |
| 5 | *T_diff_* + *T_diff_*² | 6 | 893190 | 1069 | -446589 |
| 4 | *T_diff_* | 5 | 893235 | 1115 | -446613 |
| 8 | [*T_sun_* + *T_sun_*²] *× Gs* | 12 | 895680 | 3559 | -447828 |
| 7 | *T_sun_ × Gs* | 9 | 896604 | 4483 | -448293 |
| 3 | *T_sun_* + *T_sun_*² | 6 | 896694 | 4573 | -448341 |
| 2 | *T_sun_* | 5 | 897581 | 5461 | -448786 |
| 6 | *Gs* | 6 | 913455 | 21335 | -456722 |
| 1 | 1 | 4 | 913529 | 21408 | -456760 |

Appendix 5: Table S1: Model comparison based on AICc of the model fitting the variations of the probability of being in the open when T_sun_ and T_diff_ are averaged for the last 15 minutes before the location is recorded. The models were fitted with the glmmTMB library, and included open patch availability as fixed effect in order to account for habitat availability around each location, as well as an individual and a group random effect on the intercept. The model number refers to the hypotheses and models listed in Table 1 of the main manuscript. K stands for the number of fitted parameters.

| Model # | Explicative variables | K | AICc | ΔAICc | Loglikelihood |
| --- | --- | --- | --- | --- | --- |
| 10 | [*T_diff_* + *T_diff_*²] *× Gs* | 12 | 379391 | 0 | -189683 |
| 9 | *T_diff_ × Gs* | 9 | 379466 | 76 | -189724 |
| 5 | *T_diff_* + *T_diff_*² | 6 | 379518 | 127 | -189753 |
| 4 | *T_diff_* | 5 | 379566 | 175 | -189778 |
| 8 | [*T_sun_* + *T_sun_*²] *× Gs* | 12 | 380067 | 676 | -190021 |
| 3 | *T_sun_ + T_sun_²* | 6 | 380926 | 1536 | -190457 |
| 7 | *T_sun_ × Gs* | 9 | 380944 | 1554 | -190463 |
| 2 | *T_sun_* | 5 | 380996 | 1606 | -190493 |
| 6 | *Gs* | 6 | 385213 | 5828 | -192600 |
| 1 | 1 | 4 | 385222 | 5831 | -192607 |

Appendix 5: Table S2: Model comparison based on AICc of the model fitting the variations of the probability of transition to the open when under cover when T_sun_ and T_diff_ are averaged for the last 15 minutes before the location is recorded. The models were fitted with the glmmTMB library, open patch availability at the previous location as fixed effect in order to account for habitat availability around each location,and included an individual and a group random effect on the intercept. The model number refers to the hypotheses and models listed in Table 1 of the main manuscript. K stands for the number of fitted parameters.

| Model # | Explicative variables | K | AICc | ΔAICc | Loglikelihood |
| --- | --- | --- | --- | --- | --- |
| 10 | [*T_diff_* + *T_diff_*²] *× Gs* | 12 | 320368 | 0 | -160172 |
| 9 | *T_diff_ × Gs* | 9 | 320514 | 145 | -160248 |
| 5 | *T_diff_* + *T_diff_*² | 6 | 320514 | 145 | -160251 |
| 4 | *T_diff_* | 5 | 320646 | 277 | -160318 |
| 9 | [*T_sun_* + *T_sun_*²] *× Gs* | 12 | 320904 | 536 | -160440 |
| 7 | *T_sun_ × Gs* | 9 | 320957 | 589 | -160470 |
| 3 | *T_sun_* + *T_sun_*² | 6 | 321045 | 677 | -160517 |
| 2 | *T_sun_* | 5 | 321082 | 714 | -160536 |
| 6 | *Gs* | 6 | 321366 | 998 | -160677 |
| 1 | *1* | 4 | 321368 | 1000 | -160680 |

Appendix 5: Table S3: Model comparison based on AICc of the model fitting the variations of the probability of transition under cover when in the open when T_sun_ and T_diff_ are averaged for the last 15 minutes before the location is recorded. The models were fitted with the glmmTMB library, open patch availability at the previous location as fixed effect in order to account for habitat availability around each location, and included an individual and a group random effect on the intercept. The model number refers to the hypotheses and models listed in Table 1 of the main manuscript. K stands for the number of fitted parameters.

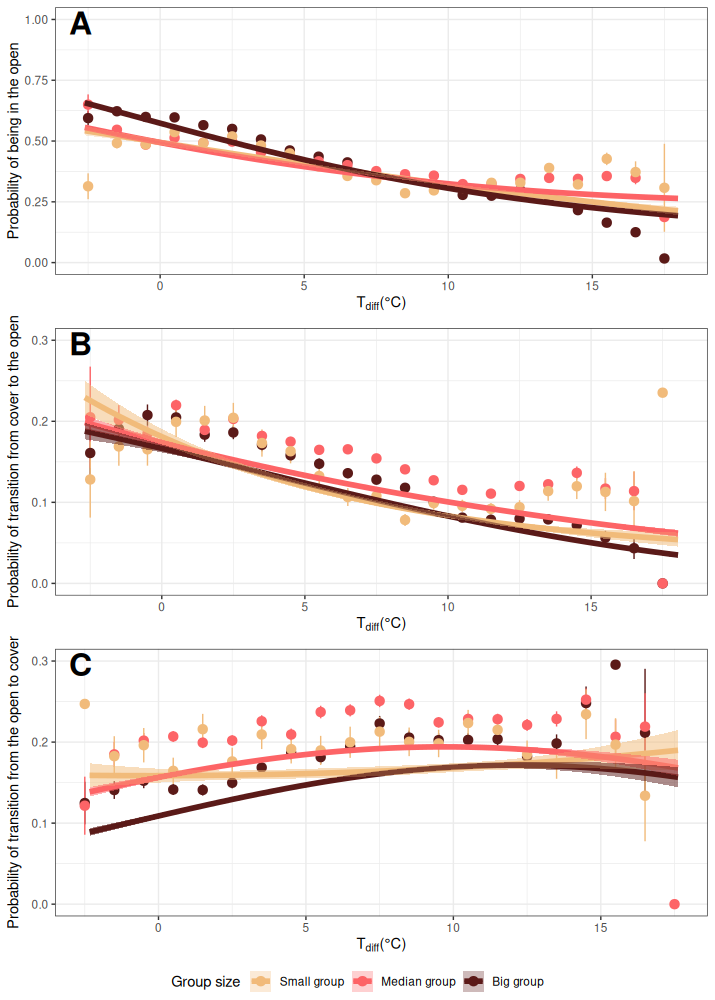
Appendix 6: Figure S1: Predictions of the best models identified by model selection for each studied behavior when T_sun_ and T_diff_ are averaged for the last 15 minutes before the location is recorded: (A) probability to be in the open vs. cover as a function of T_diff_; (B) probability to shift to the open while under cover as a function of T_diff_; (C) probability to shift to cover while in the open as a function of T_diff_. In all graphs, points are raw proportion of the behavior for each group size for bins of 1°C of T_diff_. Error bars show the calculated 95% confidence intervals around the calculated proportions. Lines are the predicted probability to do the behavior according to group size. Ribbons are the predicted 95% confidence intervals of this probability. Predictions have been made for an average open habitat availability calculated in the dataset used in the statistical model. The strong predicted effect of open habitat availability explains the deviance of the predictions to the points.

**Appendix 7 – Robustness of results of replacing instant temperature measurements to the average temperature conditions in the hour**

| Model # | Explicative variables | K | AICc | ΔAICc | Loglikelihood |
| --- | --- | --- | --- | --- | --- |
| 10 | [*T_diff_* + *T_diff_*²] *× Gs* | 12 | 895069 | 0 | -447522 |
| 9 | *T_diff_ × Gs* | 9 | 895106 | 38 | -447544 |
| 5 | *T_diff_* + *T_diff_*² | 6 | 896108 | 1040 | -448048 |
| 4 | *T_diff_* | 5 | 896122 | 1054 | -448056 |
| 8 | [*T_sun_* + *T_sun_*²] *× Gs* | 12 | 899738 | 4669 | -449857 |
| 7 | *T_sun_ × Gs* | 9 | 900231 | 5162 | -450106 |
| 3 | *T_sun_* + *T_sun_*² | 6 | 900643 | 5574 | -450316 |
| 2 | *T_sun_* | 5 | 901119 | 6051 | -450555 |
| 6 | *Gs* | 6 | 912565 | 17496 | -456276 |
| 1 | 1 | 4 | 912638 | 17569 | -456315 |

Appendix 7: Table S1: Model comparison based on AICc of the model fitting the variations of the probability of being in the open when T_sun_ and T_diff_ are averaged for the last hour before the location is recorded. The models were fitted with the glmmTMB library, and included open patch availability as fixed effect in order to account for habitat availability around each location, as well as an individual and a group random effect on the intercept. The model number refers to the hypotheses and models listed in Table 1 of the main manuscript. K stands for the number of fitted parameters.

| Model # | Explicative variables | K | AICc | ΔAICc | Loglikelihood |
| --- | --- | --- | --- | --- | --- |
| 10 | [*T_diff_* + *T_diff_*²] *× Gs* | 12 | 380096 | 0 | -190036 |
| 5 | *T_diff_* + *T_diff_*² | 6 | 380187 | 90 | -190087 |
| 9 | *T_diff_ × Gs* | 9 | 380379 | 282 | -190180 |
| 4 | *T_diff_* | 5 | 380440 | 343 | -190215 |
| 8 | [*T_sun_* + *T_sun_*²] *× Gs* | 12 | 381232 | 1135 | -190604 |
| 3 | *T_sun_ + T_sun_²* | 6 | 381291 | 1195 | -190640 |
| 7 | *T_sun_ × Gs* | 9 | 381947 | 1851 | -190965 |
| 2 | *T_sun_* | 5 | 381978 | 1882 | -190984 |
| 6 | *Gs* | 6 | 384834 | 4737 | -192411 |
| 1 | 1 | 4 | 384843 | 4746 | -192417 |

Appendix 7: Table S2: Model comparison based on AICc of the model fitting the variations of the probability of transition to the open when under cover when T_sun_ and T_diff_ are averaged for the last hour before the location is recorded. The models were fitted with the glmmTMB library, open patch availability at the previous location as fixed effect in order to account for habitat availability around each location,and included an individual and a group random effect on the intercept. The model number refers to the hypotheses and models listed in Table 1 of the main manuscript. K stands for the number of fitted parameters.

| Model # | Explicative variables | K | AICc | ΔAICc | Loglikelihood |
| --- | --- | --- | --- | --- | --- |
| 10 | [*T_diff_* + *T_diff_*²] *× Gs* | 12 | 320453 | 0 | -160215 |
| 9 | *T_diff_ × Gs* | 9 | 320567 | 113 | -160274 |
| 5 | [*T_diff_* + *T_diff_*²] | 6 | 320588 | 134 | -160288 |
| 4 | *T_diff_* | 5 | 320684 | 231 | -160337 |
| 9 | [*T_sun_* + *T_sun_*²] *× Gs* | 12 | 320714 | 261 | -160345 |
| 3 | *T_sun_* + *T_sun_*² | 6 | 320834 | 381 | -160411 |
| 7 | *T_sun_ × Gs* | 9 | 320840 | 386 | -160411 |
| 2 | *T_sun_* | 5 | 320947 | 494 | -160469 |
| 6 | *Gs* | 6 | 321037 | 584 | -160513 |
| 1 | *1* | 4 | 321039 | 585 | -160515 |

Appendix 7: Table S3: Model comparison based on AICc of the model fitting the variations of the probability of transition under cover when in the open when T_sun_ and T_diff_ are averaged for the last hour before the location is recorded. The models were fitted with the glmmTMB library, open patch availability at the previous location as fixed effect in order to account for habitat availability around each location, and included an individual and a group random effect on the intercept. The model number refers to the hypotheses and models listed in Table 1 of the main manuscript. K stands for the number of fitted parameters.

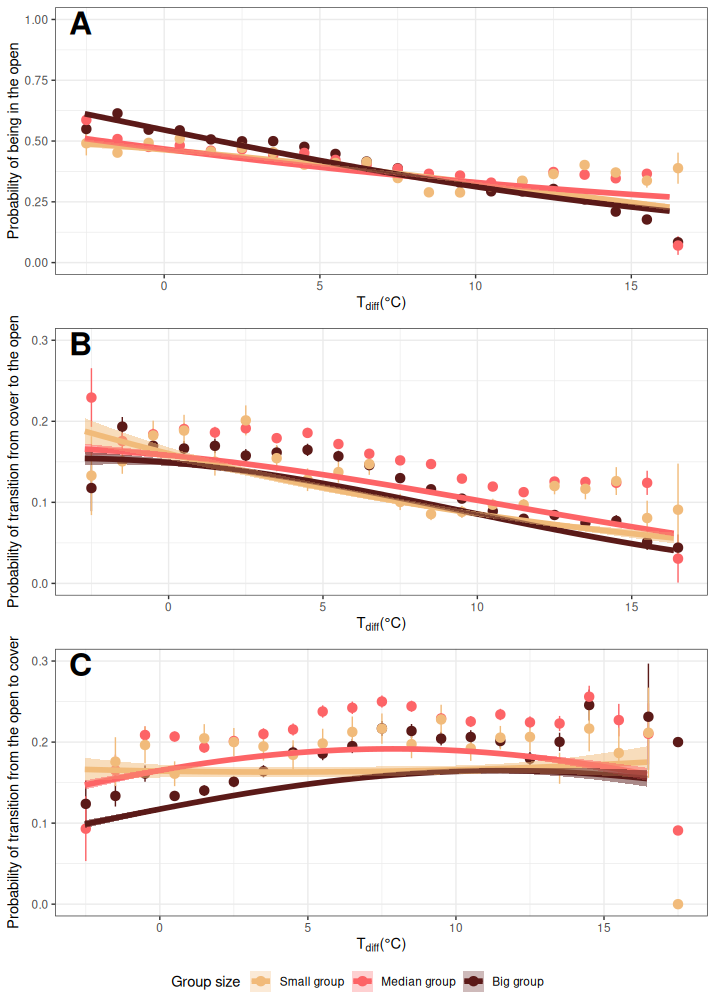
Appendix 7: Figure S1: Predictions of the best models identified by model selection for each studied behavior when T_sun_ and T_diff_ are averaged for the last hour before the location is recorded: (A) probability to be in the open vs. cover as a function of T_diff_; (B) probability to shift to the open while under cover as a function of T_diff_; (C) probability to shift to cover while in the open as a function of T_diff_. In all graphs, points are raw proportion of the behavior for each group size for bins of 1°C of T_diff_. Error bars show the calculated 95% confidence intervals around the calculated proportions. Lines are the predicted probability to do the behavior according to group size. Ribbons are the predicted 95% confidence intervals of this probability. Predictions have been made for an average open habitat availability calculated in the dataset used in the statistical model. The strong predicted effect of open habitat availability explains the deviance of the predictions to the points.
